# A day sleep promoting role of phototransduction in *Drosophila melanogaster*

**DOI:** 10.1101/2025.10.30.685496

**Authors:** Yu-Chien Hung, Mehran Akhtar, Nithish Sattoju, Xinghua Li, Steven Head, Tobias Ollerenshaw, Clelia Siefer-Gaillardin, Jashmine Arulchelvan, Ben Warren, Ko-Fan Chen

**Affiliations:** Division of Genetics and Genome Biology, University of Leicester, Leicester, UK, LE1 7RH; School of Life Sciences, Keele University, Staffordshire, UK, ST5 5BG; Magistère Européen de Génétique, University Paris Cité, Paris 75013, France

## Abstract

The daily sleep-wake cycle is a conserved behaviour defined by locomotion quiescence and enhanced responsive threshold to sensory stimuli. Both the circadian clock and sleep-homeostasis determine the daily sleep profile. Environmental light is a major sensory input and also regulates circadian clock and the balance between sleep and wakefulness. In *Drosophila*, the cellular mechanism and neural circuitry underlying light-mediated circadian synchronization are well-established, yet the direct relationship between light/visual input and sleep remains unclear. To address this knowledge gap, we measured sleep behaviour in *Drosophila* with mutations in genes involved in phototransduction and downstream neural transmission. We observed consistent day sleep fragmentation in flies with mutations in multiple phototransduction components. We also found that mutation that led to hyperpolarised *Drosophila* photoreceptors resulted in shorter day sleep. We found a severe reduction in locomotor speed in several visual mutants during normal waking time preventing assessment of their sleep-linked immobility. Taken together, our rigorous quantification of sleep in phototransduction genetic mutants reveals the key role of visual input in promoting sleep.

## 1. Introduction

Sleep is vital for good health. Disruptions to either sleep timing and/or amount has been linked to numerous metabolic and neurological diseases <u>(see reviews, Hartmann and Kempf, 2023; Wulff et al., 2010)</u>. Across the animal kingdom, sleep is defined by behavioural quiescence and increased arousal threshold to environmental stimuli such as light or mechanic vibrations. Arousal threshold within a period of sleep varies in stages that are associated with specific electrophysiological signatures in electroencephalogram (EEG) and local field potentials (LFPs) in the brain (Lakhiani et al., 2023). With its versatile genetic toolkit and conserved behavioural and electrophysiological features of sleep (Jagannathan et al., 2024; Shafer and Keene, 2021), the fruit fly *Drosophila melanogaster* has been used for identifying the biological basis and the conserved function of sleep. In parallel to murine investigations, studies in *Drosophila* have demonstrated that neuronal redox status may form a cellular basis of sleep (Hartmann and Kempf, 2023) and sleep stages play roles in synaptic scaling that is important for learning and memory (Dissel et al., 2015; Gilestro et al., 2009; Tainton-Heap et al., 2021; Yap et al., 2017). Recent investigations in *Drosophila* have also shown that sleep is required for effective maintenance of the immune system, gut microbiome, and waste clearance (Alphen et al., 2021; Vaccaro et al., 2020; Vincent et al., 2022).

Daily sleep onset and duration are regulated by two intrinsic mechanisms: the circadian clock and sleep homeostasis. The circadian clock defines the baseline sleep-wake level across 24 hours, whereas sleep homeostasis maintains the overall level of sleep amount. In *Drosophila*, the neural circuits that generate circadian rhythm are composed of 242 clock neurons (Reinhard et al., 2024) that express the core molecular clock mechanism. Whereas the homeostatic control of sleep is distributed across the following neural domains: Pars Intercerebralis (PI), Mushroom Body (MB), Fan-Shaped Body (FSB) and Ellipsoid Body (EB) (Jones et al., 2025; Shafer and Keene, 2021). The circadian clock and sleep homeostasis are likely coordinated, as clock neurons modulate the FSB-mediated sleep homeostat (Hsu et al., 2025).

Light is a prominent environmental stimulus that regulates both circadian clock and sleep homeostasis to maintain an animal’s temporal niche and sufficient sleep. The synchronisation between environmental light regime and the neural activity of clock neurons is called circadian photoentrainment. Visual inputs are part of photoentrainment in *Drosophila*. The visual system in the fly is composed of peripheral light sensing neurons: photoreceptors in compound eyes, HB-eyelet and ocelli. Photoreceptors in the eye are subdivided into outer achromatic R1-R6 and inner chromatic R7/R8 neurons, both of which sends their synaptic outputs to downstream visual processing brain areas called the optic lobes and ventrolateral neuropils (Nériec and Desplan, 2016). Crucially, light signals are converted into depolarisation of the neuronal membrane via a cellular process called phototransduction in the photoreceptors of the compound eye and HB-eyelet. Phototransduction requires the light-sensing GPCR Rhodopsins 1-6 (Rh1-6) and their downstream signal cascades that ultimately activate cation channels depolarising the photoreceptors (Hardie, 2012). Activation of photoreceptors results in synaptic release of histamine or acetylcholine to communicate with clock neurons either directly or indirectly via a neural network in the optic lobes (Alejevski et al., 2019; Ogueta et al., 2020, 2018; Rieger et al., 2003; Schlichting et al., 2016; Tang et al., 2022; Xiao et al., 2023). Apart from via the visual system, *Drosophila* circadian clocks are entrained by cell autonomous light-sensing mechanisms mediated by *cryptochrome* (*cry*)*, quasimodo* (*qsm*) or *Rhodopsin 7* (*Rh7*) (Chen et al., 2011; Ni et al., 2017; Peschel et al., 2009).

Light can directly regulate sleep by either promoting or supressing sleep via different neural pathways in the murine system (Altimus et al., 2008; Pilorz et al., 2016). As a crepuscular species, *Drosophila* sleep both in the day (light) and in the night (darkness). While current evidence indicates that distinct neurogenetic networks may control day and night sleep (Chen et al., 2019; Damulewicz et al., 2020; Hsu et al., 2025; Ishimoto et al., 2012; Lamaze et al., 2018; Mazzotta et al., 2020; Sengupta et al., 2019), the mechanism underlying light-mediated regulation of the *Drosophila* sleep homeostat is far from clear. Previous studies indicate that midday sleep (i.e., siesta during six hours of the midday) can be enhanced by high-intensity light via the HB-eyelet-mediated cholinergic pathway (Rieger et al., 2007; Schlichting et al., 2019). Separate studies however, have shown contradictory findings for the role of compound eye photoreceptors in sleep regulation: synaptic inhibition in these photoreceptors promote sleep (Damulewicz et al., 2020), yet the optogenetic activation of the same photoreceptors in another investigation did not result in sleep suppression (Kirszenblat et al., 2019). Moreover, artificial activation of the direct downstream target of eye photoreceptors, lamina monopolar neuron 2 (L2 LMC), resulted in drastic sleep loss (Muraro and Ceriani, 2015). This finding contradicts the increased sleep observed in flies with reduced histamine (Oh et al., 2013), which is the main neurotransmitters released from photoreceptors to inhibit L2 LMC neurons (Pantazis et al., 2008). Furthermore, visual experience in light was recently found to promote night sleep in *Drosophila* (Kirszenblat et al., 2019), suggesting light and/or vision *per se* serve as sleep homeostat drivers, similar to in vertebrates (Horne and Walmsley, 1976; Korf et al., 2017; Miyamoto et al., 2003).

A recent systematic study of mutants with defects in various light input pathways indicate that only *Rh7* is required for shade sleep preference in the day, but the study did not investigate the baseline sleep level in these mutants (Meyerhof et al., 2024). We therefore investigated the role of light input pathways in *Drosophila* sleep by focusing on the role of the visual system. Using the conventional infrared beam-based sleep assay (Chen et al., 2019; Chiu et al., 2010), we surveyed baseline sleep profiles in three groups of flies: strains with mutations in genes involved in (i) phototransduction and (ii) downstream histaminergic neurotransmission (Hardie, 2012), and (iii) those with artificial inhibition of photoreceptors. We also performed electroretinogram (ERG) examine the phototransduction/neurotransmission defects in these above mutants. By carefully examining sleep and ERG parameters, we identified a novel role of visual phototransduction in driving sleep.

## 2. Materials and methods

### 2.1 Fly husbandry

*Drosophila* strains were kept in standard fly food (glucose 6.93% w/v, agar 0.5% w/v, cornmeal, 6.3% w/v, yeast, 4.38% w/v, propionic acid, 0.26% v/v, 0.2% v/v nipagin) at 25°C 12-hour light dark cycle. Long-term fly stocks are kept in the same food and light regime at 18°C. The Canton S and Oregon R wildtype strains were obtained from the Kyriacou Lab (University of Leicester, UK), while *w^iso31^* (i.e. *iso*) control was obtained from the Jepson lab (University College London, UK). The following visual mutants, Gal4 and UAS transgenic flies were ordered from Bloomington Drosophila Stock Center (Indiana University, USA) (stock number): *trp^1^* (5692); *trp^MB^* (23636); *trpl^302^* (31433); *trpl^MB^* (29134); *ninaE^8^* (2001); *ninaE^5^* (3531); *ninaE^17^* (5701); *norpA^P24^* (9048); *hdc^MB^* (25260); *hdc^JK910^* (64203); *ort^1^* (1133); *ort^attP^* (84559); *Rh1-Gal4* (8691); *gmr-Gal4* (1104); *UAS-kir2* (6595); *UAS-TeTxLC* (28837); *UAS-myr-mRFP* (7118). All Gal4 and UAS transgenic flies were brought into the *w^iso31^* background via five generation outcrossing following *[w+]* eye colour marker cassette. For *Rh1-Gal4* outcrossing, we generated *UAS-myr-mRFP/+; Rh1-Gal4/+* flies which expresses RFP in the eyes, we then crossed this line to *w^iso31^* and used this RFP signal as a visible marker under a fluorescent stereoscope to ensure outcrossing offspring still contained both *UAS-myr-mRFP* and *Rh1-Gal4* transgenes. The final stock was then established by balancer crossing with *w^iso31C^;Sco/CyO; MKRS/TM6B* and then *w^iso31^;+/+; MKRS/TM6B* for removing *UAS-myr-mRFP*. Among other lines, *hdc^MB^*, *ort^attP^*, *trp^MB^*, and *trpl^MB^* contained either GFP or RFP visible markers, but the rest are classic missense mutations, making outcrossing without laborious PCR-based sequencing difficult. For simplicity, sleep experiments for all non-Gal4/UAS mutants were conducted without outcrossing and the only consistent results among mutant alleles of each gene were interpreted. Following the unexpected but consistent night sleep among *hdc* mutants, we decided to minimise the genetic background difference among these lines: since the precise genetic lesion of *hdc^JK910^* on the *hdc* loci at the 2^nd^ chromosome is unknown, only the 1^st^ and 3^rd^ chromosomes of *hdc^JK910^* were replaced with those from *w^iso31^* stock via the following successive crossings with balancer lines (gifts from Jepson lab):

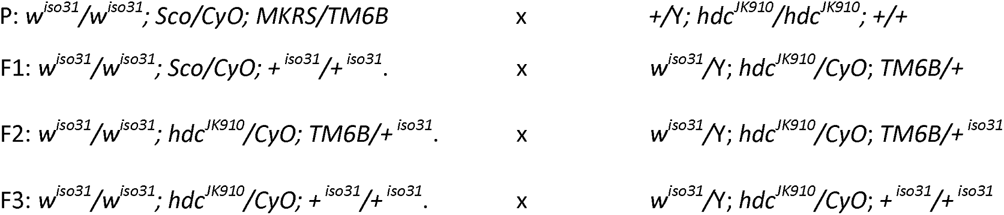

Separately, *hdc^MB^* flies were brought into the *w^iso31^* background via five generation outcrossing following the *GFP* eye marker.

### 2.2 Sleep recording and analysis

*Drosophila* sleep, particularly day time sleep, is highly dependent on by sex, mating and nutrient status (Dove et al., 2017; Duhart et al., 2023; Isaac et al., 2009; Wu et al., 2018). Typically, mated females can sleep for as little as 150-200 minutes compared to the 300-400 minutes shown by males (out of the total 720 minutes of day time). To capture both sleep increase and decrease upon genetic manipulation, we focused our investigation on virgin male flies. Moreover, by focusing on males, our data is comparable to the previous decades of photoentrainment investigations of *Drosophila* circadian research that has been conducted mainly on male visual mutants (e.g. Rieger et al., 2003 and see review in Mazzotta et al., 2020). As such, we would caution against extending our findings to those of sleep in female flies.

Prior to the sleep experiment, *Drosophila* strains or crossings were reared in food with a richer recipe (wheat germ 1% w/v, soy 0.8% w/v, glucose/sucrose 3.3%/1.5% w/v, treacle 3% v/v, agar 1% w/v, cornmeal 1.5% w/v, yeast 3.5% w/v, propionic acid 0.5% v/v, nipagin 0.1% v/v), so as to keep the rearing conditions consistent with previous experimental conditions (Chen et al., 2019, 2011; Lamaze et al., 2018). We also note that in general, all strains of flies are less motile or less reproductively successful in standard fly food (as described above in 2.1). Sleep recordings were performed as described in (Chen et al., 2019). Three to four day-old virgin adult males were transferred to typical behaviour tubes containing food (2% w/v agar and 4% w/v sucrose at one end and cotton wool at the other), and housed in typical Drosophila Activity Monitor (DAM, Trikinetics Inc.) for 3 days of 12hours light-dark cycle at 25°C in a temperature and light controlled incubator (MIR-254-PE, PHCbi). The lighting source was a day light fluorescent lamp (FL15D, 6500K) with intensity between 1.2 and 6.7 w/m^2^ depending on the distance to the lamp. With conservative conversion with luminance surface area between 0.25 to 1 m^2^, our light intensity was approximately equal to 402 to 1608 lux. The locomotion of each fly was recorded as number of infrared beam crosses per minute. The locomotion data for day 3 was transferred to a previously customised Excel Calculator (Chen et al., 2019) to calculate all sleep episodes (defined as any period of immobility ≥ 5 minutes) along 24 hours. Additionally, the following parameters were calculated for each fly: **overall sleep profile**: minutes of sleep per 30 mins across 24 hours; **day sleep**: total minutes of sleep during 12 hours of day; **night sleep**: total minutes of sleep during 12 hours of night; **day average sleep bout length**: day sleep minutes divided by total number of sleep bout during 12 hours of day; **night average sleep bout length:** night sleep minutes divided by total number of sleep bout during 12 hours of night; **day waking activity**: number of beam crosses divided by total wake minutes during 12 hours of day**; night waking activity**: number of beam crosses divided by total wake minutes during 12 hours of night.

### 2.3 Electroretinogram (ERG)

Three-to-five day-old adult virgin male flies were inserted into a p1000 plastic pipette tip, so that its head protruded from the narrow ending. The head was fixed using melted honey wax applied between the thorax and head without blocking the mouthpart and eyes. The pipette tip mounted fly was kept in the dark for 20 mins and then transferred into the ERG recording rig. Glass electrodes (Borosilicate) with resistance of ∼5 MO and filled with 3 M KCl were inserted into the compound eye (recording) and head (reference). Each fly was left in the dark for 3 minutes before being exposed to four light intensities: 0.004, 0.02, 0.1 and 0.5 w/m^2^ at 540nm LED green light source (M530L4-C1, ThorLab). These intensities fall within the liner range of receptor potential response. We stimulated with 1000 ms exposure to each light intensity and repeated the procedure two further times to obtain an average value of three presentations at each light intensity for each fly. The potential difference between the electrodes was amplified 10x, and acquired by Neuroprobe Amplifier model 1600 and 8-channel data acquisition interface DA-8 (A-M systems) using the SciWorks Data Acquisition & Analysis Suite.

### 2.4 Data display and statistics

The `ggplot2` package (3.5.2) in R (4.4.2) used to display sleep and ERG data. X-Y plots were used to display overall sleep profiles for each compared genotype. Boxplots with average values of each individual were used to display sleep parameters data. We calculated the average voltage change of on transient, receptor potential and off transient in the ERG profiles of each replicate for each fly (Figure 1H). The value for each fly was then plotted against light intensity on logarithmic scales by `ggplot2`. Considering the variability in sleep data (Chen et al., 2019; Lamaze et al., 2018), statistical significance was identified by Kruskal-Wallis rank sum test for non-parametric multiple comparison (`kruskal.test`). Pairwise Wilcoxon rank sum exact test with multiple comparison Benjamini-Hochberg correction (`pairwise.wilcox.test`) was used to specifically identify difference between pairwise genotypes. Unlike sleep data, ERG voltage changes data were pooled across four intensities when identifying genotypical difference via Pairwise Wilcoxon rank sum exact test. Significance level is set at adj.p<0.05. Effect size *r* is calculated by `wilcox_effsize` with magnitude categories: small; moderate and large assigned. Importantly, while Benjamini-Hochberg correction is required to correct the p-values, the effect size *r* does not change by multiple comparison, as it is defined by *Z/√n*. All statistic comparisons, p values, effect size and sample size are listed in Table1 to Table 6 for the corresponding Figure 1 to Figure 5.

**Figure 1.**
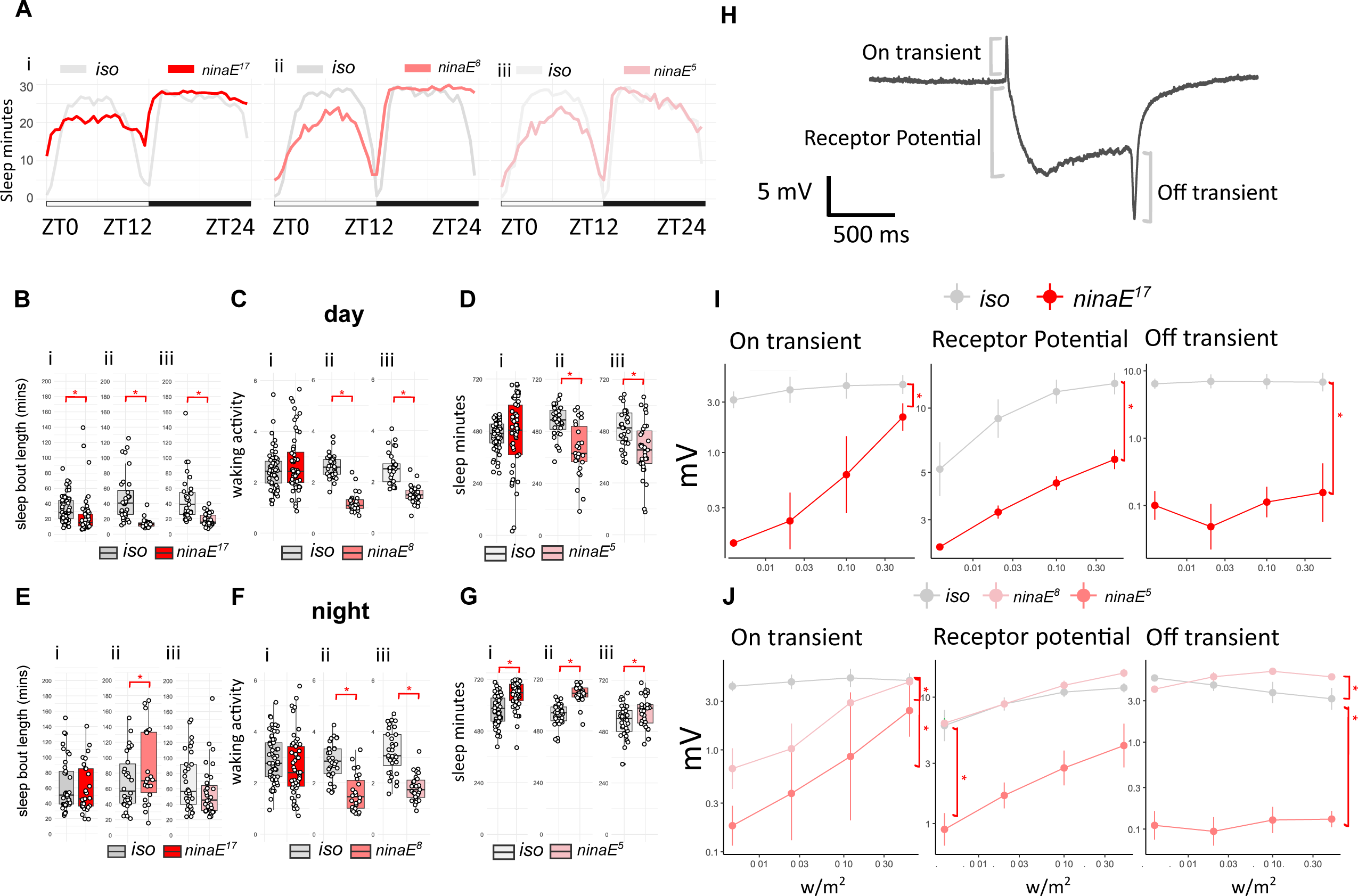
sleep profile, parameters and ERG quantification for *ninaE* mutants. **A.** Sleep traces of average sleep minutes per 30 minutes over 12 hours light-dark cycle. ZT0: light on, ZT12: light off, ZT24: end of night. Sleep traces of *iso* control and *ninaE* mutants are colour coded in grey and red shades (i-iii). **B.** Average day sleep bout length (minutes), **C.** Average day wake activity (beam breaks per waking minutes), **D.** Average day sleep (minutes), E. Average night sleep bout length (minutes), **F.** Average night wake activity (beam breaks per waking minutes), and **G.** Average night sleep (minutes) for *ninaE^17^* (i), *ninaE^8^* (ii) and *ninaE^5^* (iii) compared to iso controls. Average values for each individual are plotted as filled circles except for those outside scales. **H.** An example ERG profile from iso control with 1000 ms green light flash that evokes voltage changes of on-transient, receptor potential and off-transient. **I.** Mean and standard deviation for voltage change (mV) in on transient, receptor potential and off-transient of ERG upon four light intensity (0.004, 0.02, 0.1 and 0.5 w/m^2^) flash for *ninaE^17^* and controls. **J.** Mean and standard deviation for voltage change (mV) in on transient, receptor potential and off-transient of ERG upon four light intensities (0.004, 0.02, 0.1 and 0.5 w/m^2^) 1000ms flash for *ninaE^5^*, *ninaE^8^* and controls. Plot scales are in either liner (A-G) and logarithmic (I-J). Asterisk indicates statistical significance (p<0.05) between mutants and iso controls by pairwise Wilcoxon rank sum exact test without (B-H) and with multiple comparison, Benjamini-Hochberg correction (I, followed by Kruskal-Wallis rank sum test). See **Table 1** for sample size (n=26-60 for sleep; n=3-6 for ERG, with 4 replicates for each light intensity), p-values and effect size for each comparison.

## 3. Result

### 3.1 Defects in phototransduction genes caused short day sleep bout

To verify the role of visual phototransduction genes in sleep regulation, we tested eight mutant fly strains. These strains are mutants of the following four genes: *Rhodopsin 1/ninaE*, *Phospholipase C/norpA*, and the two cation channels *trp* and t*rpl*. These included both classic missense mutations, and when available, more recently characterised mutations using P-element insertion. To analyse the details of sleep-wake profiles under a typical day of 12 hours light-12 hours dark, we examined three conventional parameters for both day and night period: total sleep, average sleep bout length and waking activity.

#### 3.1.1 ninaE

Upon exposure to light, 11-cis 3-hydroxy-retinal, the chromophore in the Rhodopsin protein undergoes photoisomerization, transforming Rhodopsin to Metarhodopsin which then triggers a downstream GPCR signal cascade. Rhodopsin 1/ninaE protein is the main light-sensing GPCR in the outer photoreceptors R1-R6 of the *Drosophila* compound eye. Reduction or loss of ninaE abolishes or reduces photoisomerization and therefore blocks the light-induced photoreceptor potential and transients that are typically measured for R1-R6 in ERG (Hardie, 2012).

Many visual tasks, such as navigation, phototaxis, optomotor and escape, require R1-R6 activity. We therefore first tested the sleep profile of the class null allele *ninaE^17^* (O’Tousa et al., 1985). During the day, *ninaE^17^* mutants show a lower midday sleep peak in the sleep profile (Figure 1Ai) but the amount of total sleep is comparable to the white eye control, *w^iso31^* (here after *iso*, Figure 1Di). However, we found that the average sleep bout length in *ninaE^17^* is shorter than the control, indicating sleep fragmentation (Figure 1Bi). During the night, we observed increased sleep amount (Figure 1Gi) while the average sleep bout length is similar to the control (Figure 1Ei). Since sleep measurements in *Drosophila* depend on locomotion, an increase in sleep can be falsely claimed due to defects in locomotion. Therefore, “waking activity”, i.e., locomotor activity per waking minutes, was designed as a conventional measure to detangle such a compound scenario (Andretic and Shaw, 2005; Harbison and Sehgal, 2008). We applied this measurement and found that waking activity is similar between *ninaE^17^* and the control in both day and night (Figure 1Ci,Fi).

*ninaE^17^* flies contain a secondary *ebony^1^* mutation that impacts sleep-wake cycles (Rossi et al., 2015; Suh and Jackson, 2007). We therefore tested two other *ninaE* hypomorphs: *ninaE^5^* and *ninaE^8^* (Scavarda et al., 1983). Both mutants show similar midday sleep peak reductions to those seen in *ninaE^17^* flies (Figure 1Aii-iii). This observation is consistent with the significant reduced day sleep observed in either mutant when compared to the control (Figure 1Dii-iii). Similarly, both mutants also show shorter average sleep bout length in the day (Figure 1Bii-iii). Intriguingly, the loss of sleep in the two *ninaE* hypomorphs was observed despite of the loss of waking activity in the day (Figure 1Cii-iii). During the night, *ninaE* hypomorphs both show increased sleep (Figure 1Gii-iii), but only *ninaE^8^* mutant shows significant increase of the average sleep bout length compared to control flies (Figure 1Eii-iii). However, mutants severely reduce their waking activity (Figure 1Fii-iii with effect size *r* between 0.67 and 0.82, Table 1). Therefore, the increase in night sleep observed here cannot be certain, and may simply reflect the loss of locomotor activity. To confirm *ninaE* mutants harbour phototransduction defects, we measured light-evoked receptor potentials through Electroretinograms (ERG, Figure 1H). Consistent with previous reports, *ninaE^17^* and *ninaE^8^* exhibit reduced light induction of receptor potential, on and off transients (Figure 1I-J), whereas *ninaE^5^* retains certain function (see normal receptor potential and off-transient in *ninaE^5^*, Figure 1J). Taken together, our data indicates that the reduction, or loss of, *Drosophila* Rhodopsin 1 (ninaE) protein results in the reduction of day sleep bout length, and that phototransduction is required to drive day sleep pressure. However, recent studies show support for a non-visual function of ninaE (Zanini et al., 2018). To verify if ninaE protein alone or the phototransduction in general regulates day sleep bout length, we further examined the sleep-regulatory roles for three other key proteins for phototransduction.

**Table 1.**
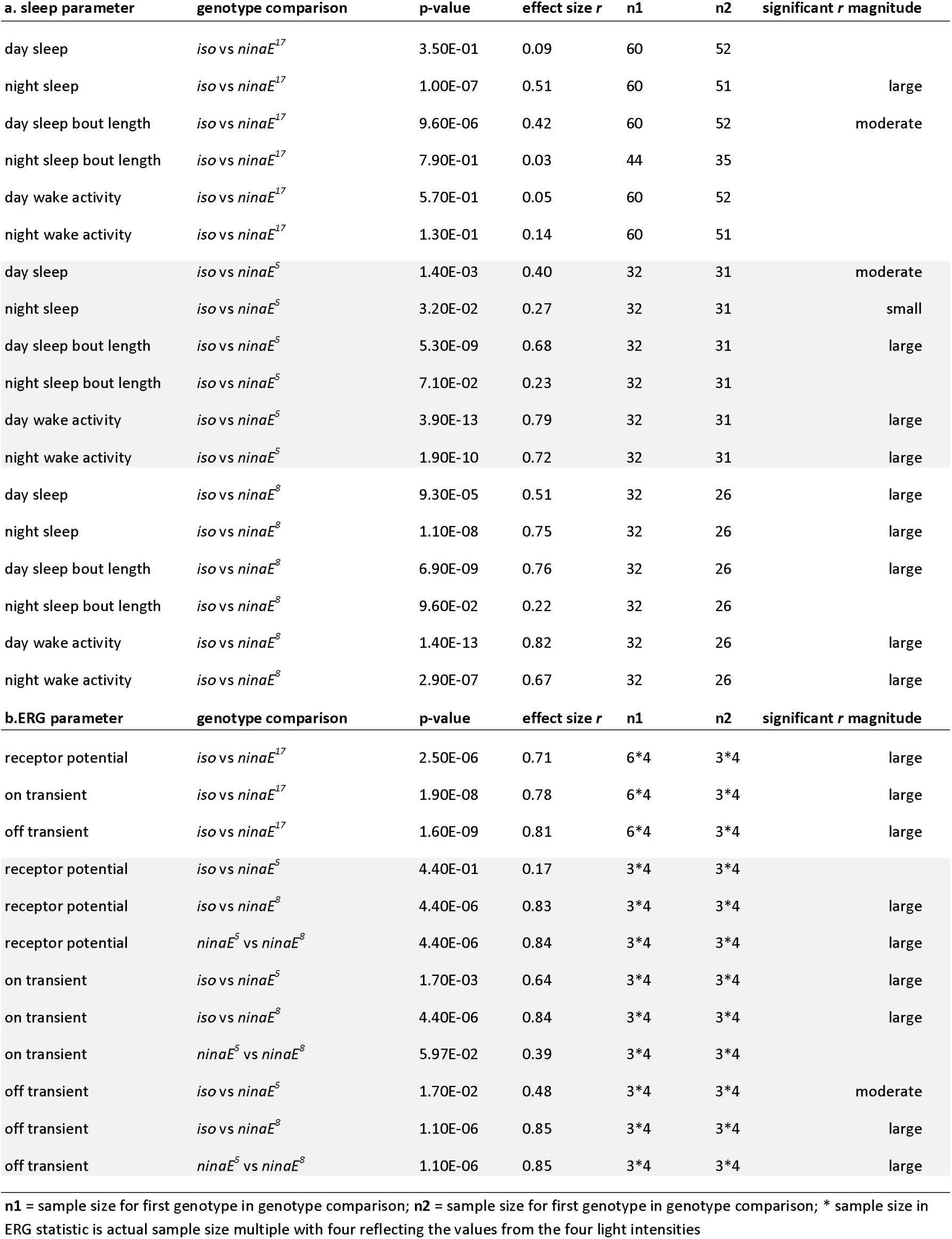
Statistics of a. sleep and b. ERG parameters for *ninaE* mutants.

| a. sleep parameter | genotype comparison | p-value | effect size *r* | n1 | n2 | significant *r* magnitude |
| --- | --- | --- | --- | --- | --- | --- |
| day sleep | *iso* vs *ninaE*<sup>17</sup> | 3.50E-01 | 0.09 | 60 | 52 |  |
| night sleep | *iso* vs *ninaE*<sup>17</sup> | 1.00E-07 | 0.51 | 60 | 51 | large |
| day sleep bout length | *iso* vs *ninaE*<sup>17</sup> | 9.60E-06 | 0.42 | 60 | 52 | moderate |
| night sleep bout length | *iso* vs *ninaE*<sup>17</sup> | 7.90E-01 | 0.03 | 44 | 35 |  |
| day wake activity | *iso* vs *ninaE*<sup>17</sup> | 5.70E-01 | 0.05 | 60 | 52 |  |
| night wake activity | *iso* vs *ninaE*<sup>17</sup> | 1.30E-01 | 0.14 | 60 | 51 |  |
| day sleep | *iso* vs *ninaE*<sup>5</sup> | 1.40E-03 | 0.40 | 32 | 31 | moderate |
| night sleep | *iso* vs *ninaE*<sup>5</sup> | 3.20E-02 | 0.27 | 32 | 31 | small |
| day sleep bout length | *iso* vs *ninaE*<sup>5</sup> | 5.30E-09 | 0.68 | 32 | 31 | large |
| night sleep bout length | *iso* vs *ninaE*<sup>5</sup> | 7.10E-02 | 0.23 | 32 | 31 |  |
| day wake activity | *iso* vs *ninaE*<sup>5</sup> | 3.90E-13 | 0.79 | 32 | 31 | large |
| night wake activity | *iso* vs *ninaE*<sup>5</sup> | 1.90E-10 | 0.72 | 32 | 31 | large |
| day sleep | *iso* vs *ninaE*<sup>8</sup> | 9.30E-05 | 0.51 | 32 | 26 | large |
| night sleep | *iso* vs *ninaE*<sup>8</sup> | 1.10E-08 | 0.75 | 32 | 26 | large |
| day sleep bout length | *iso* vs *ninaE*<sup>8</sup> | 6.90E-09 | 0.76 | 32 | 26 | large |
| night sleep bout length | *iso* vs *ninaE*<sup>8</sup> | 9.60E-02 | 0.22 | 32 | 26 |  |
| day wake activity | *iso* vs *ninaE*<sup>8</sup> | 1.40E-13 | 0.82 | 32 | 26 | large |
| night wake activity | *iso* vs *ninaE*<sup>8</sup> | 2.90E-07 | 0.67 | 32 | 26 | large |
| b. ERG parameter | genotype comparison | p-value | effect size *r* | n1 | n2 | significant *r* magnitude |
| receptor potential | *iso* vs *ninaE*<sup>17</sup> | 2.50E-06 | 0.71 | 6*4 | 3*4 | large |
| on transient | *iso* vs *ninaE*<sup>17</sup> | 1.90E-08 | 0.78 | 6*4 | 3*4 | large |
| off transient | *iso* vs *ninaE*<sup>17</sup> | 1.60E-09 | 0.81 | 6*4 | 3*4 | large |
| receptor potential | *iso* vs *ninaE*<sup>5</sup> | 4.40E-01 | 0.17 | 3*4 | 3*4 |  |
| receptor potential | *iso* vs *ninaE*<sup>8</sup> | 4.40E-06 | 0.83 | 3*4 | 3*4 | large |
| receptor potential | *ninaE*<sup>5</sup> vs *ninaE*<sup>8</sup> | 4.40E-06 | 0.84 | 3*4 | 3*4 | large |
| on transient | *iso* vs *ninaE*<sup>5</sup> | 1.70E-03 | 0.64 | 3*4 | 3*4 | large |
| on transient | *iso* vs *ninaE*<sup>8</sup> | 4.40E-06 | 0.84 | 3*4 | 3*4 | large |
| on transient | *ninaE*<sup>5</sup> vs *ninaE*<sup>8</sup> | 5.97E-02 | 0.39 | 3*4 | 3*4 |  |
| off transient | *iso* vs *ninaE*<sup>5</sup> | 1.70E-02 | 0.48 | 3*4 | 3*4 | moderate |
| off transient | *iso* vs *ninaE*<sup>8</sup> | 1.10E-06 | 0.85 | 3*4 | 3*4 | large |
| off transient | *ninaE*<sup>5</sup> vs *ninaE*<sup>8</sup> | 1.10E-06 | 0.85 | 3*4 | 3*4 | large |

n1 = sample size for first genotype in genotype comparison; n2 = sample size for first genotype in genotype comparison; \* sample size in ERG statistic is actual sample size multiple with four reflecting the values from the four light intensities

#### 3.1.2 norpA

Metarhodopsin triggers Gqα activation and enzymatic reaction of Phospholipase C (PLC) that produces secondary messengers. The norpA protein is the main PLC in the visual system, and the classic mutant *norpA^p24^* exhibits severe visual defects (Gao et al., 2008). The sleep profile of *norpA^p24^* shows decline of day sleep from the midday and a secondary dip at midnight compared to the control (Figure 2A). There is also a phase advance of night sleep in *norpA^p24^* mutants as compared to the controls. Such an advance was also noticeable in locomotor profile of another *norpA* allele in a previous study (Ogueta et al., 2018). The changes in sleep profile coincides with the reduction in day sleep (Figure 2D) and shorter bout length in both day and night (Figure 2B, 2E). Similar to *ninaE^5^* and *ninaE^8^*, the *norpA^p24^* mutant has low waking activity (Figure 2C, 2F, effect size 0.39/0.75, Table 2). We also confirmed abnormal ERG profiles in *norpA^p24^* (Figure 2H).

**Figure 2.**
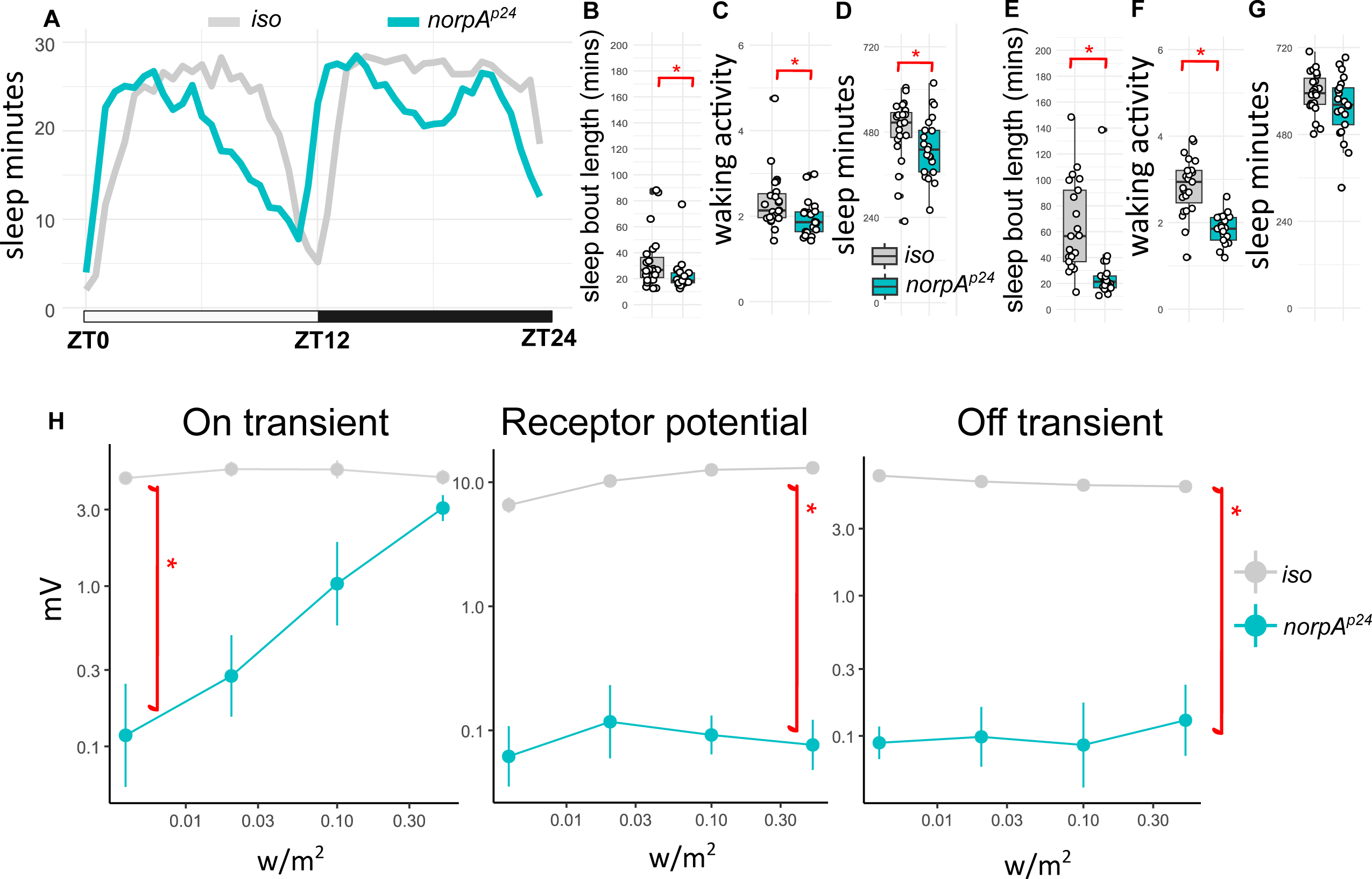
sleep profile, parameters and ERG quantification for *norpA* mutants. **A.** Sleep traces of average sleep minutes per 30 minutes over 12 hours light-dark cycle. ZT0: light on, ZT12: light off, ZT24: end of night. Sleep traces for *iso* control and *norpA^p24^* mutants are colour coded in grey and cyan. **B.** Average day sleep bout length (minutes), **C.** Average day wake activity (beam breaks per waking minutes), **E.** Average night sleep bout length (minutes), **F.** Average night wake activity (beam breaks per waking minutes), **G.** Average night sleep (minutes) for *norpA^p24^* compared to iso controls. Average values for each individual are plotted as filled circles except for those outside scales (B-G). **H.** mean and standard deviation for voltage change (mV) in on transient, receptor potential and off-transient of ERG upon four light intensity (0.004, 0.02, 0.1 and 0.5 w/m^2^) flash for *norpA^p24^* and *iso* controls. Plot scales are in either liner (A-G) and logarithmic (H). Asterisk indicates statistical significance (p<0.05) between mutants and iso controls by pairwise Wilcoxon rank sum exact test. See **Table 2** for sample size (n=21-23 for sleep; n=4 for ERG, with 4 replicates for each light intensity), p-values and effect size for each comparison.

**Table 2.** Statistics of a. sleep and b. ERG parameters for *norpA* mutants.

| a. sleep parameter | genotype comparison | p-value | effect size *r* | n1 | n2 | significant *r* magnitude |
| --- | --- | --- | --- | --- | --- | --- |
| day sleep bout length | *iso* vs *norpA*<sup>p24</sup> | 9.40E-03 | 0.39 | 23 | 21 | moderate |
| night sleep bout length | *iso* vs *norpA*<sup>p24</sup> | 2.50E-06 | 0.66 | 23 | 21 | large |
| day sleep | *iso* vs *norpA*<sup>p24</sup> | 3.20E-02 | 0.32 | 23 | 21 | moderate |
| night sleep | *iso* vs *norpA*<sup>p24</sup> | 1.20E-01 | 0.23 | 23 | 21 |  |
| day wake activity | *iso* vs *norpA*<sup>p24</sup> | 9.40E-03 | 0.39 | 23 | 21 | moderate |
| night wake activity | *iso* vs *norpA*<sup>p24</sup> | 1.70E-07 | 0.72 | 23 | 21 | large |
| b.ERG parameter | genotype comparison | p-value | effect size *r* | n1 | n2 | significant *r* magnitude |
| receptor potential | *iso* vs *norpA*<sup>p24</sup> | 1.50E-06 | 0.85 | 4*4 | 4*4 | large |
| on transient | *iso* vs *norpA*<sup>p24</sup> | 3.30E-09 | 0.85 | 4*4 | 4*4 | large |
| off transient | *iso* vs *norpA*<sup>p24</sup> | 1.50E-06 | 0.85 | 4*4 | 4*4 | large |

n1 = sample size for first genotype in genotype comparison; n2 = sample size for first genotype in genotype comparison; \* sample size in ERG statistic is actual sample size multiple with four reflecting the values from the four light intensities

#### 3.1.3 trp and trpl

Activation of the two ion channels, trp and trpl in the last step of phototransduction causes the crucial cation influx resulting in depolarisation of the photoreceptor. We therefore examined the sleep profile of four mutant lines, two P-element insertion mediated mutants: *trp^MB^*, *trpl^MB^* (Chen et al., 2015) and two classic null mutants: *trp^1^*, and *trpl^302^* (Cosens and Manning, 1969; Leung et al., 2000; Niemeyer et al., 1996). Except for *trp^1^*, these lines have white eyes, like the *ninaE* and *norpA* mutants as described above.

The *trp^1^* mutant was first isolated from a red eye wild type strain, but the precise background was not known (Cosens and Manning, 1969), so we compared its sleep profile to two widely used wild type strains in sleep and vision research: *Canton S* and *Oregon R.* Significant reductions in day and night sleep and average sleep bout length are shown in *trp^1^* mutants as compared to either wild type flies (Figure 3Ai, 3Bi and 3Ei, effect size: 0.29-0.54, Table 3). The waking activity of *trp^1^* is comparable to *Canton S* but higher than *Oregon R*, and has limited impact to sleep loss (Figure 3Ci and 3Fi). The sleep profile of *trp^MB^* is distinct from *trp^1^, norpA* or *ninaE* mutants, with day sleep level similar to controls but a significant night sleep loss (Figure 3Aii, 3Gii). There is no significant change in sleep bout length detected in *trp^MB^* flies (Figure 3Aii and 3Eii). Although, *trp^MB^* flies sleep a similar amount during the day, compared to the control, their waking activity is significantly higher (Figure 3Cii).

**Figure 3.**
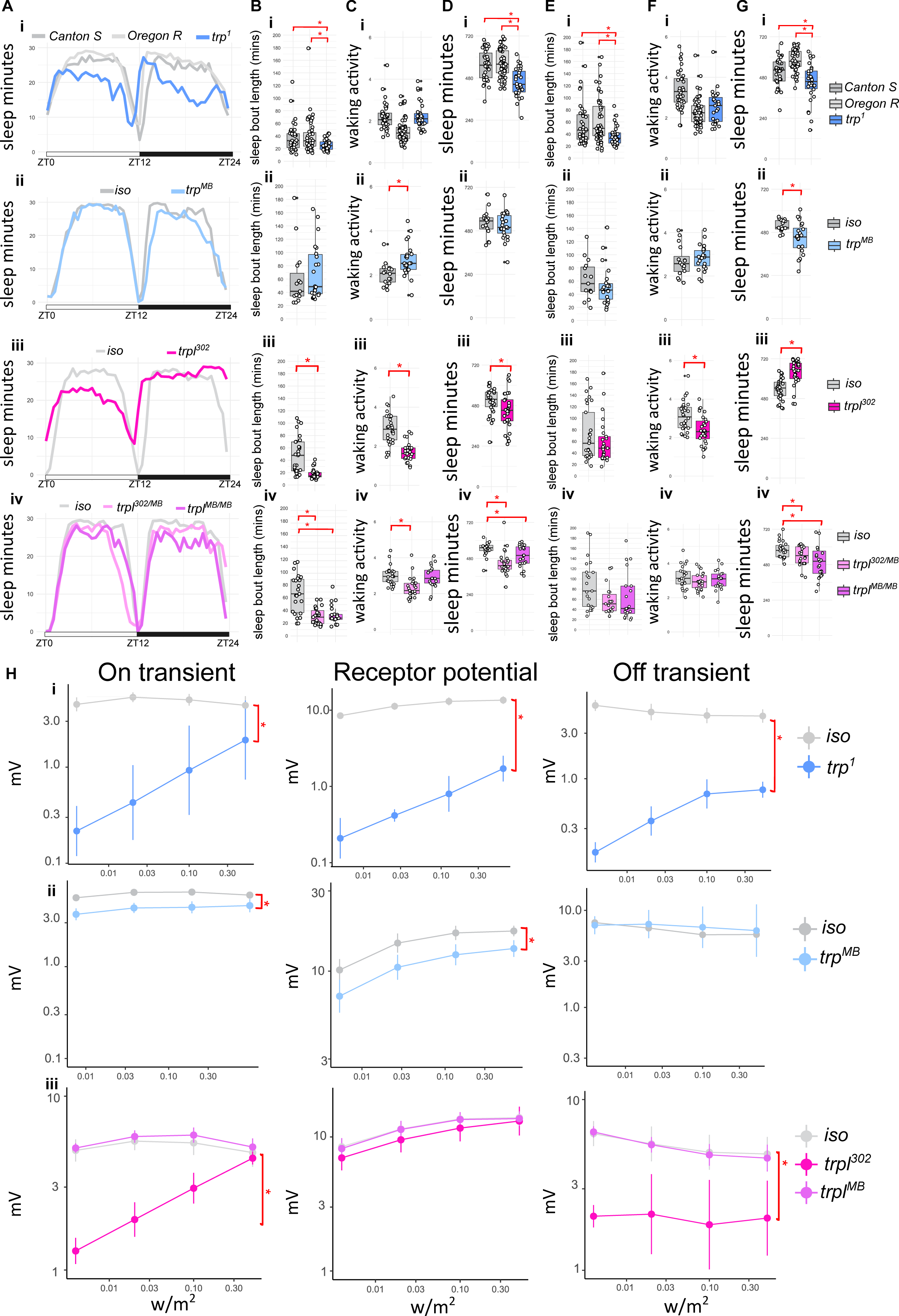
sleep profile, parameters and ERG quantification for *trp* and *trpl* mutants. **A.** Sleep traces of average sleep minutes per 30 minutes over 12 hours light-dark cycle. ZT0: light on, ZT12: light off, ZT24: end of night. Sleep traces for *Canton S* (i), *Oregon R* (i) and *iso* controls (ii-iv) are coloured coded in the shades of greys; *trp* (i-ii) and *trpl* (iii-iv) mutants are colour coded in shades of blue and purples. **B.** Average day sleep bout length (minutes), **C.** Average day wake activity (beam breaks per waking minutes), **D.** Average day sleep (minutes), **E.** Average night sleep bout length (minutes), **F.** Average night wake activity (beam breaks per waking minutes), **G.** Average night sleep (minutes) for *trp^1^* (i), *trp^MB^* (ii), *trpl^302^* (iii) and *trpl^302/MB^* or *trpl^MB^* (iv) compared to controls. Average values for each individual are plotted as filled circles except for those outside scales. **H.** Mean and standard deviation for voltage change (mV) in on transient, receptor potential and off-transient of ERG upon four light intensity (0.004, 0.02, 0.1 and 0.5 w/m^2^) flash for *trp^1^* (i), *trp^MB^* (ii), *trpl^302^* (iii) and *trpl^MB^* (iii) compared to controls. Plot scales are in either liner (A-G) and logarithmic (H). Asterisk indicates statistical significance (p<0.05) between mutants and controls by pairwise Wilcoxon rank sum exact test without (Aii-Gii, Aiii-Giii, Hi and Hii) and with multiple comparison, Benjamini-Hochberg correction (Ai-Gi, Aiv-Giv, and Hiii followed by Kruskal-Wallis rank sum test). See **Table 3 and Table 4** for sample size (n=16-53 for sleep; n=3 for ERG, with 4 replicates for each light intensity), p-values and effect size for each comparison.

**Table 3.** Statistics of a. sleep and b. ERG parameters for *trp* mutants.

| a. sleep parameter | genotype comparison | p-value | effect size *r* | n1 | n2 | significant *r* magnitude |
| --- | --- | --- | --- | --- | --- | --- |
| day sleep | *Canton S* vs *Oregon R* | 5.33E-01 | 0.07 | 38 | 53 |  |
| day sleep | *Canton S* vs *trp*<sup>1</sup> | 1.50E-04 | 0.47 | 38 | 30 | moderate |
| day sleep | *Oregon R* vs *trp*<sup>1</sup> | 2.90E-06 | 0.54 | 53 | 30 | large |
| night sleep | *Canton S* vs *Oregon R* | 1.20E-04 | 0.41 | 38 | 53 | moderate |
| night sleep | *Canton S* vs *trp*<sup>1</sup> | 1.63E-02 | 0.29 | 38 | 30 | small |
| night sleep | *Oregon R* vs *trp*<sup>1</sup> | 2.30E-07 | 0.59 | 53 | 30 | large |
| day sleep bout length | *Canton S* vs *Oregon R* | 8.66E-01 | 0.02 | 38 | 53 |  |
| day sleep bout length | *Canton S* vs *trp*<sup>1</sup> | 1.40E-02 | 0.32 | 38 | 30 | moderate |
| day sleep bout length | *Oregon R* vs *trp*<sup>1</sup> | 1.20E-02 | 0.32 | 53 | 30 | moderate |
| night sleep bout length | *Canton S* vs *Oregon R* | 6.61E-01 | 0.05 | 38 | 53 |  |
| night sleep bout length | *Canton S* vs *trp*<sup>1</sup> | 5.20E-03 | 0.35 | 38 | 30 | moderate |
| night sleep bout length | *Oregon R* vs *trp*<sup>1</sup> | 1.20E-03 | 0.39 | 53 | 30 | moderate |
| day wake activity | *Canton S* vs *Oregon R* | 8.20E-08 | 0.58 | 38 | 53 | large |
| day wake activity | *Canton S* vs *trp*<sup>1</sup> | 7.70E-01 | 0.04 | 38 | 30 |  |
| day wake activity | *Oregon R* vs *trp*<sup>1</sup> | 2.90E-07 | 0.57 | 53 | 30 | large |
| night wake activity | *Canton S* vs *Oregon R* | 1.20E-07 | 0.58 | 38 | 53 | large |
| night wake activity | *Canton S* vs *trp*<sup>1</sup> | 1.60E-02 | 0.31 | 38 | 30 | moderate |
| night wake activity | *Oregon R* vs *trp*<sup>1</sup> | 6.70E-02 | 0.20 | 53 | 30 |  |
| day sleep | *iso* vs *trp*<sup>MB</sup> | 5.80E-01 | 0.09 | 16 | 24 |  |
| night sleep | *iso* vs *trp*<sup>MB</sup> | 3.70E-03 | 0.46 | 16 | 24 | moderate |
| day sleep bout length | *iso* vs *trp*<sup>MB</sup> | 3.20E-01 | 0.16 | 16 | 24 |  |
| night sleep bout length | *iso* vs *trp*<sup>MB</sup> | 3.90E-01 | 0.14 | 16 | 24 |  |
| day wake activity | *iso* vs *trp*<sup>MB</sup> | 1.40E-03 | 0.49 | 16 | 24 | moderate |
| night wake activity | *iso* vs *trp*<sup>MB</sup> | 2.90E-01 | 0.17 | 16 | 24 |  |
| b.ERG parameter | genotype comparison | p-value | effect size *r* | n1 | n2 | significant *r* magnitude |
| receptor potential | *iso* vs *trp*<sup>1</sup> | 7.40E-07 | 0.85 | 3*4 | 3*4 | large |
| on transient | *iso* vs *trp*<sup>1</sup> | 2.20E-05 | 0.78 | 3*4 | 3*4 | large |
| off transient | *iso* vs *trp*<sup>1</sup> | 7.40E-07 | 0.85 | 3*4 | 3*4 | large |
| receptor potential | *iso* vs *trp*<sup>MB</sup> | 4.40E-03 | 0.53 | 3*4 | 4*4 | large |
| on transient | *iso* vs *trp*<sup>MB</sup> | 2.60E-07 | 0.83 | 3*4 | 4*4 | large |
| off transient | *iso* vs *trp*<sup>MB</sup> | 5.70E-01 | 0.11 | 3*4 | 4*4 |  |

n1 = sample size for first genotype in genotype comparison; n2 = sample size for first genotype in genotype comparison; \* sample size in ERG statistic is actual sample size multiple with four reflecting the values from the four light intensities

This relative normal sleep-wake profile may relate to the less severe reduction in receptor potential and light on/off transients in the ERG of *trp^MB^* as compared to those of the *trp^1^* null mutant (Figure 3Hi and 3Hii, effect size *r* = 0.11-0.83 vs 0.78-0.85, Table 3).

On the other hand, both *trpl^302^* and *trpl^MB^* mutants exhibit day sleep profiles similar to *ninaE* or *norpA* mutants (Figure 3Aiii and 3Aiv), with significant reduced day sleep and average bout length (Figure 3Biii, 3Biv, 3Diii and 3Div). Night sleep profiles and waking activities for the two *trpl* mutants diverged: *trpl^302^* sleep more whereas *trpl^MB^* sleep less (Figure 3Giii, 3Giv). No significant changes in night sleep bout length were observed in both mutants as compared to the controls (Figure 3Eiii and 3Eiv). *trpl^302^* mutants generally have low waking activity, whereas *trpl^MB^* maintains relative normal waking activity as compared to the controls (Figure 3Ciii, 3Civ, 3Fiii and 3Fiv). *trpl^302^* mutant is in *cn* and *bw* mutation background, which may affect the sleep. We therefore tested *w; trpl^302/MB^* flies to reduce the impact of *cn bw* mutation. Similar to *trpl^302^* and *trpl^MB^*, the *w; trpl^302/MB^* transheterozygote mutant showed shorter day sleep and sleep bout length (Figure 3Biv). As seen in previous reports (Chen et al., 2015; Leung et al., 2000) both mutants have limited impact on ERG profiles, *trpl^302^* flies show a large reduction of on/off transient (effect size 0.73/0.81, Table 4), whereas *trpl^MB^* mutants maintain normal ERGs, possibly due to synaptic compensatory mechanism (Figure 3Hiii). Nevertheless, our data indicate the reduced trpl function and expression level cause sleep loss in the day.

**Table 4.** Statistics of a. sleep and b. ERG parameters for *trpl* mutants.

| a. sleep parameter | genotype comparison | p-value | effect size *r* | n1 | n2 | significant *r* magnitude |
| --- | --- | --- | --- | --- | --- | --- |
| day sleep | *iso* vs *trpl*<sup>302</sup> | 1.40E-02 | 0.32 | 30 | 30 | moderate |
| night sleep | *iso* vs *trpl*<sup>302</sup> | 1.40E-06 | 0.62 | 30 | 30 | large |
| day sleep bout length | *iso* vs *trpl*<sup>302</sup> | 1.40E-06 | 0.59 | 30 | 30 | large |
| night sleep bout length | *iso* vs *trpl*<sup>302</sup> | 6.90E-01 | 0.05 | 30 | 28 |  |
| day wake activity | *iso* vs *trpl*<sup>302</sup> | 1.10E-08 | 0.68 | 30 | 30 | large |
| night wake activity | *iso* vs *trpl*<sup>302</sup> | 9.70E-04 | 0.43 | 30 | 28 | moderate |
| day sleep | *iso/iso* vs *trpl*<sup>302</sup>/*trpl*<sup>MB</sup> | 3.70E-05 | 0.63 | 23 | 24 | large |
| day sleep | *iso/iso* vs *trpl*<sup>MB</sup>/*trpl*<sup>MB</sup> | 3.71E-02 | 0.32 | 23 | 21 | moderate |
| day sleep | *trpl*<sup>302</sup>/*trpl*<sup>MB</sup> vs *trpl*<sup>MB</sup>/*trpl*<sup>MB</sup> | 2.57E-02 | 0.35 | 24 | 21 | moderate |
| night sleep | *iso/iso* vs *trpl*<sup>302</sup>/*trpl*<sup>MB</sup> | 7.32E-02 | 0.28 | 24 | 22 |  |
| night sleep | *iso/iso* vs *trpl*<sup>MB</sup>/*trpl*<sup>MB</sup> | 1.56E-02 | 0.39 | 24 | 23 | moderate |
| night sleep | *trpl*<sup>302</sup>/*trpl*<sup>MB</sup> vs *trpl*<sup>MB</sup>/*trpl*<sup>MB</sup> | 1.52E-01 | 0.22 | 22 | 23 |  |
| day sleep bout length | *iso/iso* vs *trpl*<sup>302</sup>/*trpl*<sup>MB</sup> | 1.60E-04 | 0.59 | 23 | 23 | large |
| day sleep bout length | *iso/iso* vs *trpl*<sup>MB</sup>/*trpl*<sup>MB</sup> | 1.60E-04 | 0.58 | 23 | 21 | large |
| day sleep bout length | *trpl*<sup>302</sup>/*trpl*<sup>MB</sup> vs *trpl*<sup>MB</sup>/*trpl*<sup>MB</sup> | 8.69E-01 | 0.03 | 23 | 21 |  |
| night sleep bout length | *iso/iso* vs *trpl*<sup>302</sup>/*trpl*<sup>MB</sup> | 2.20E-01 | 0.23 | 23 | 22 |  |
| night sleep bout length | *iso/iso* vs *trpl*<sup>MB</sup>/*trpl*<sup>MB</sup> | 8.42E-02 | 0.31 | 23 | 21 |  |
| night sleep bout length | *trpl*<sup>302</sup>/*trpl*<sup>MB</sup> vs *trpl*<sup>MB</sup>/*trpl*<sup>MB</sup> | 4.82E-01 | 0.13 | 22 | 21 |  |
| day wake activity | *iso/iso* vs *trpl*<sup>302</sup>/*trpl*<sup>MB</sup> | 1.20E-04 | 0.58 | 23 | 23 | large |
| day wake activity | *iso/iso* vs *trpl*<sup>MB</sup>/*trpl*<sup>MB</sup> | 9.63E-01 | 0.01 | 23 | 21 |  |
| day wake activity | *trpl*<sup>302</sup>/*trpl*<sup>MB</sup> vs *trpl*<sup>MB</sup>/*trpl*<sup>MB</sup> | 1.04E-03 | 0.50 | 23 | 21 | large |
| night wake activity | *iso/iso* vs *trpl*<sup>302</sup>/*trpl*<sup>MB</sup> | 2.91E-01 | 0.21 | 23 | 22 |  |
| night wake activity | *iso/iso* vs *trpl*<sup>MB</sup>/*trpl*<sup>MB</sup> | 6.31E-01 | 0.12 | 23 | 21 |  |
| night wake activity | *trpl*<sup>302</sup>/*trpl*<sup>MB</sup> vs *trpl*<sup>MB</sup>/*trpl*<sup>MB</sup> | 7.10E-01 | 0.07 | 22 | 21 |  |
| b.ERG parameter | genotype comparison | p-value | effect size *r* | n1 | n2 | significant *r* magnitude |
| receptor potential | *iso* vs *trpl*<sup>302</sup> | 2.20E-01 | 0.26 | 6*4 | 3*4 |  |
| receptor potential | *iso* vs *trpl*<sup>MB</sup> | 8.60E-01 | 0.03 | 6*4 | 5*4 |  |
| receptor potential | *trpl*<sup>302</sup> vs *trpl*<sup>MB</sup> | 2.20E-01 | 0.26 | 3*4 | 5*4 |  |
| on transient | *iso* vs *trpl*<sup>302</sup> | 2.40E-06 | 0.73 | 6*4 | 3*4 | large |
| on transient | *iso* vs *trpl*<sup>MB</sup> | 1.50E-01 | 0.22 | 6*4 | 5*4 |  |
| on transient | *trpl*<sup>302</sup> vs *trpl*<sup>MB</sup> | 1.00E-05 | 0.80 | 3*4 | 5*4 | large |
| off transient | *iso* vs *trpl*<sup>302</sup> | 4.80E-09 | 0.81 | 6*4 | 3*4 | large |
| off transient | *iso* vs *trpl*<sup>MB</sup> | 9.90E-01 | 0.00 | 6*4 | 5*4 |  |
| off transient | *trpl*<sup>302</sup> vs *trpl*<sup>MB</sup> | 5.30E-08 | 0.81 | 3*4 | 5*4 | large |

n1 = sample size for first genotype in genotype comparison; n2 = sample size for first genotype in genotype comparison; \* sample size in ERG statistic is actual sample size multiple with four reflecting the values from the four light intensities

**Table 4.** Statistics of a. sleep and b. ERG parameters for *hdc* mutants.

| a. sleep parameter | genotype comparison | p-value | effect size *r* | n1 | n2 | significant *r* magnitude |
| --- | --- | --- | --- | --- | --- | --- |
| day sleep | *hdc*<sup>MB</sup> vs *iso* | 1.10E-01 | 0.29 | 16 | 16 |  |
| day sleep | *hdc*<sup>JK910</sup> vs *iso* | 7.30E-05 | 0.60 | 24 | 23 | large |
| night sleep | *hdc*<sup>MB</sup> vs *iso* | 1.40E-02 | 0.44 | 16 | 16 | moderate |
| night sleep | *hdc*<sup>JK910</sup> vs *iso* | 4.99E-03 | 0.42 | 24 | 23 | moderate |
| day sleep bout length | *hdc*<sup>MB</sup> vs *iso* | 4.50E-01 | 0.14 | 16 | 16 |  |
| day sleep bout length | *hdc*<sup>JK910</sup> vs *iso* | 1.08E-01 | 0.25 | 24 | 23 |  |
| night sleep bout length | *hdc*<sup>MB</sup> vs *iso* | 1.50E-02 | 0.43 | 16 | 16 | moderate |
| night sleep bout length | *hdc*<sup>JK910</sup> vs *iso* | 2.50E-06 | 0.66 | 24 | 23 | large |
| day wake activity | *hdc*<sup>MB</sup> vs *iso* | 8.70E-01 | 0.03 | 16 | 16 |  |
| day wake activity | *hdc*<sup>JK910</sup> vs *iso* | 2.50E-01 | 0.19 | 24 | 23 |  |
| night wake activity | *hdc*<sup>MB</sup> vs *iso* | 5.60E-01 | 0.11 | 16 | 16 |  |
| night wake activity | *hdc*<sup>JK910</sup> vs *iso* | 1.50E-06 | 0.68 | 24 | 23 | large |
| b. ERG parameter | genotype comparison | p-value | effect size *r* | n1 | n2 | significant *r* magnitude |
| receptor potential | *iso* vs *hdc*<sup>JK910</sup> | 8.00E-01 | 0.24 | 3*4 | 3*4 |  |
| receptor potential | *iso* vs *hdc*<sup>MB</sup> | 4.00E-01 | 0.06 | 3*4 | 3*4 |  |
| receptor potential | *hdc*<sup>JK910</sup> vs *hdc*<sup>MB</sup> | 3.80E-01 | 0.32 | 3*4 | 3*4 |  |
| on transient | *iso* vs *hdc*<sup>JK910</sup> | 4.20E-05 | 0.72 | 3*4 | 3*4 | large |
| on transient | *iso* vs *hdc*<sup>MB</sup> | 2.20E-04 | 0.79 | 3*4 | 3*4 | large |
| on transient | *hdc*<sup>JK910</sup> vs *hdc*<sup>MB</sup> | 9.32E-01 | 0.02 | 3*4 | 3*4 |  |
| off transient | *iso* vs *hdc*<sup>JK910</sup> | 5.50E-05 | 0.85 | 3*4 | 3*4 | large |
| off transient | *iso* vs *hdc*<sup>MB</sup> | 2.20E-06 | 0.85 | 3*4 | 3*4 | large |
| off transient | *hdc*<sup>JK910</sup> vs *hdc*<sup>MB</sup> | 4.80E-04 | 0.72 | 3*4 | 3*4 | large |

n1 = sample size for first genotype in genotype comparison; n2 = sample size for first genotype in genotype comparison; \* sample size in ERG statistic is actual sample size multiple with four reflecting the values from the four light intensities

### 3.2 Divergent sleep function in histaminergic pathway

*Drosophila* photoreceptors are divided into Rh1-expressing (R1-R6) and non-Rh1-exppressing neurons (R7/R8). The R1-R6 photoreceptors form synapses onto lamina monopolar neurons (LMC), while the R7/R8 send their output to the medulla in the optic lobes. Histamine is the primary neurotransmitter for Rh1-expessing photoreceptors. Therefore, we tested if mutants with defects in histamine synthesis or postsynaptic reception show altered sleep. Histidine decarboxylase (*hdc*) is the main histamine synthesis enzyme (Pantazis et al., 2008). We investigated the sleep profiles for the two *hdc* mutants (*hdc^JK910^* and *hdc^MB^*) that were showed to have reduced histamine levels in the photoreceptors (Burg et al., 1993; Stenesen et al., 2015).

As expected, both mutants selectively reduce on/off transients’ levels, the components in ERG that reflect synaptic transmission between photoreceptors and LMCs (Figure 4Hi). Surprisingly, significant sleep defects were detected in the night instead of during the day. During the day, increased day sleep and bout length are found in *hdc^JK910^* mutants as compared to the controls, whereas *hdc^MB^* flies show comparable day sleep and bout length to the controls (Figure 4Bi-ii and 4Di-ii). The waking activities in the day are similar between *hdc* mutants and controls (Figure 4Ci-ii). During the night, both *hdc* mutants sleep less with shorter sleep bouts (Figure 4Ei-ii and 4Gi-ii). Only *hdc^JK910^* mutants have reduced waking activities compared to the controls at night (Figure 4Fi-ii). Two histamine receptors, ort and HisCl1, are detected in *Drosophila* CNS, and they are both histamine-gated chloride channels (Pantazis et al., 2008). Transcriptomic data in *Drosophila* visual system suggests that ort is expressed selectively in neurons postsynaptic to photoreceptors, whereas HisCl1 is expressed almost exclusively in photoreceptors as auto-receptors (Davis et al., 2020). We therefore verified the sleep profile in ort mutants to investigate the role of histamine signal downstream to photoreceptors.

**Figure 4.**
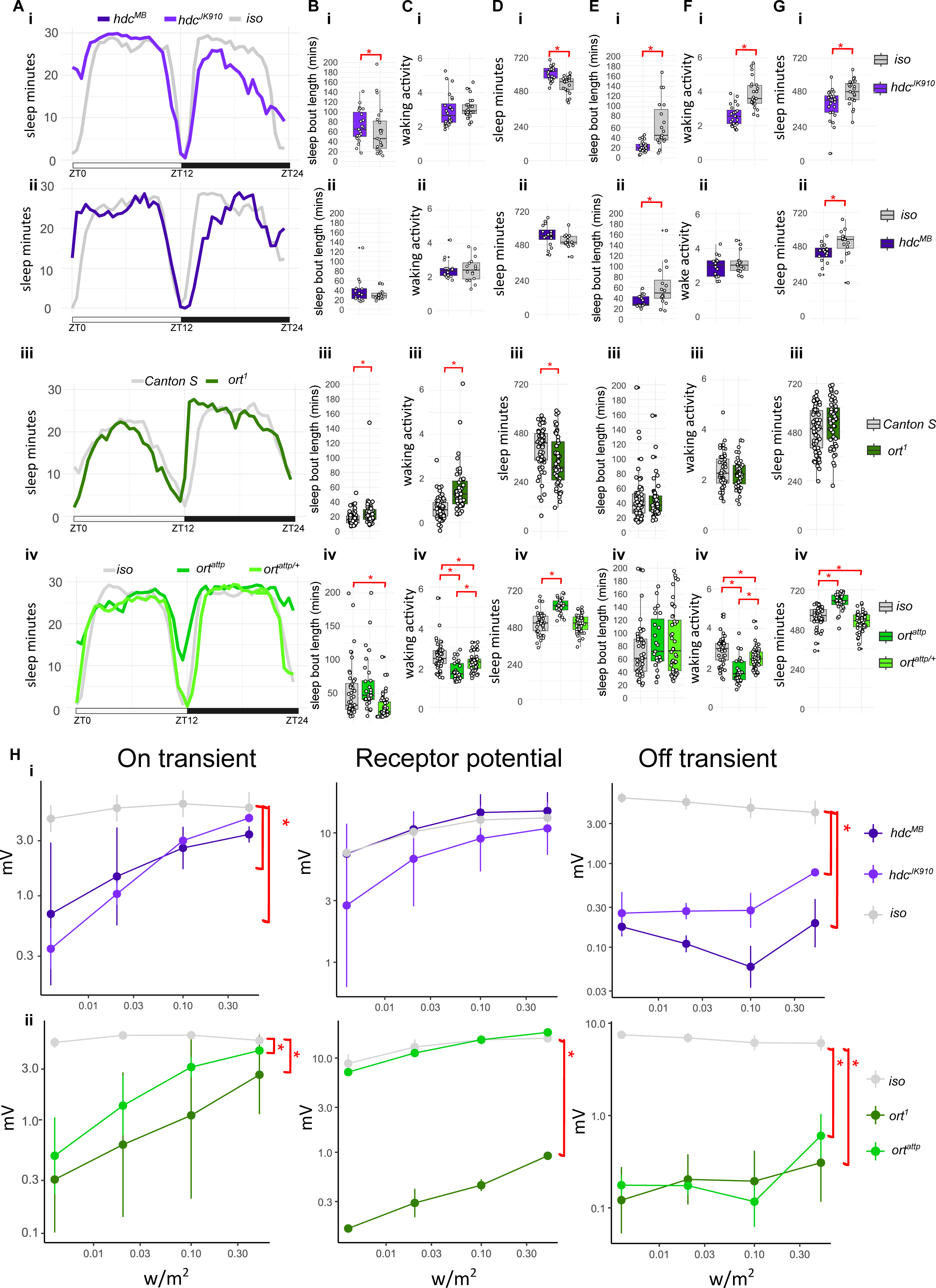
sleep profile, parameters and ERG quantification for *hdc* and *ort* mutants. **A.** Sleep traces of average sleep minutes per 30 minutes over 12 hours light-dark cycle. ZT0: light on, ZT12: light off, ZT24: end of night. Sleep traces for *iso* (i-ii, iv) and *Canton S* (iii) controls are coloured coded in the shades of greys; *hdc* (i-ii) and *ort* (iii-iv) mutants are colour coded in shades of indigos and greens. **B.** Average day sleep bout length (minutes), **C.** Average day wake activity (beam breaks per waking minutes), **D.** Average day sleep (minutes), **E.** Average night sleep bout length (minutes), **F.** Average night wake activity (beam breaks per waking minutes), **G.** Average night sleep (minutes) for *hdc* mutants (i-ii), *ort^1^* and *ort^attp/+^* or *ort^attp^* (iii-iv) compared to controls. Average values for each individual are plotted as filled circles except for those outside scales (B-G). **H.** Mean and standard deviation for voltage change (mV) in on transient, receptor potential and off-transient of ERG upon four light intensity (0.004, 0.02, 0.1 and 0.5 w/m^2^) flash for *hdc* (i) and *ort* (ii) mutants compared to controls. Plot scales are in either liner (A-G) and logarithmic (H). Asterisk indicates statistical significance (p<0.05) between mutants and controls by pairwise Wilcoxon rank sum exact test without (Aii-Gii) and with multiple comparison, Benjamini-Hochberg correction (Ai-Hi, Aiii-Giii, and Hii followed by Kruskal-Wallis rank sum test). See **Table 5** for sample size (n=29-60 for sleep; n=3-5 for ERG, with 4 replicates for each light intensity), p-values and effect size for each comparison.

First, we monitored the sleep profile of the classic *ort^1^* mutant against *Canton S*, as they share the same genetic backgrounds (Inoue et al., 1989). A small but significant day time sleep loss is detected in *ort^1^* flies (Figure 4Dii, effective size 0.27, Table 5). *ort^1^* mutants sleep for longer bouts during the day but have higher day waking activity as compared to *Canton S* (Figure 4Biii and 4Ciii). In contrast to *hdc* mutants, no night sleep phenotype is detected in *ort^1^* flies (Figure 4Eiii-4Giii). We also tested a recent CRISPR-mediated null mutation *ort^attp^* in the white eye background (Deng et al., 2019). The sleep profile of *ort^attp^* is distinct from *ort^1^*, showing more sleep as compared to control (Figure 4Div and 4Giv). However, the waking activities of *ort^attp^* are severely reduced (Figure 4Civ and 4Fiv, effective size 0.55/0.60, Table 5) therefore compounding the observed sleep phenotype. Intriguingly, shorter day sleep bout length and a small reduction of night sleep are detected in *ort^attp/+^* heterozygotes (Figure 4Biv and 4Giv), accompanying improved waking activity (Figure 4Civ and 4Fiv, wake activity *ort^attp^* vs *ort^attp/+^*, Table 5).

**Table 5.**
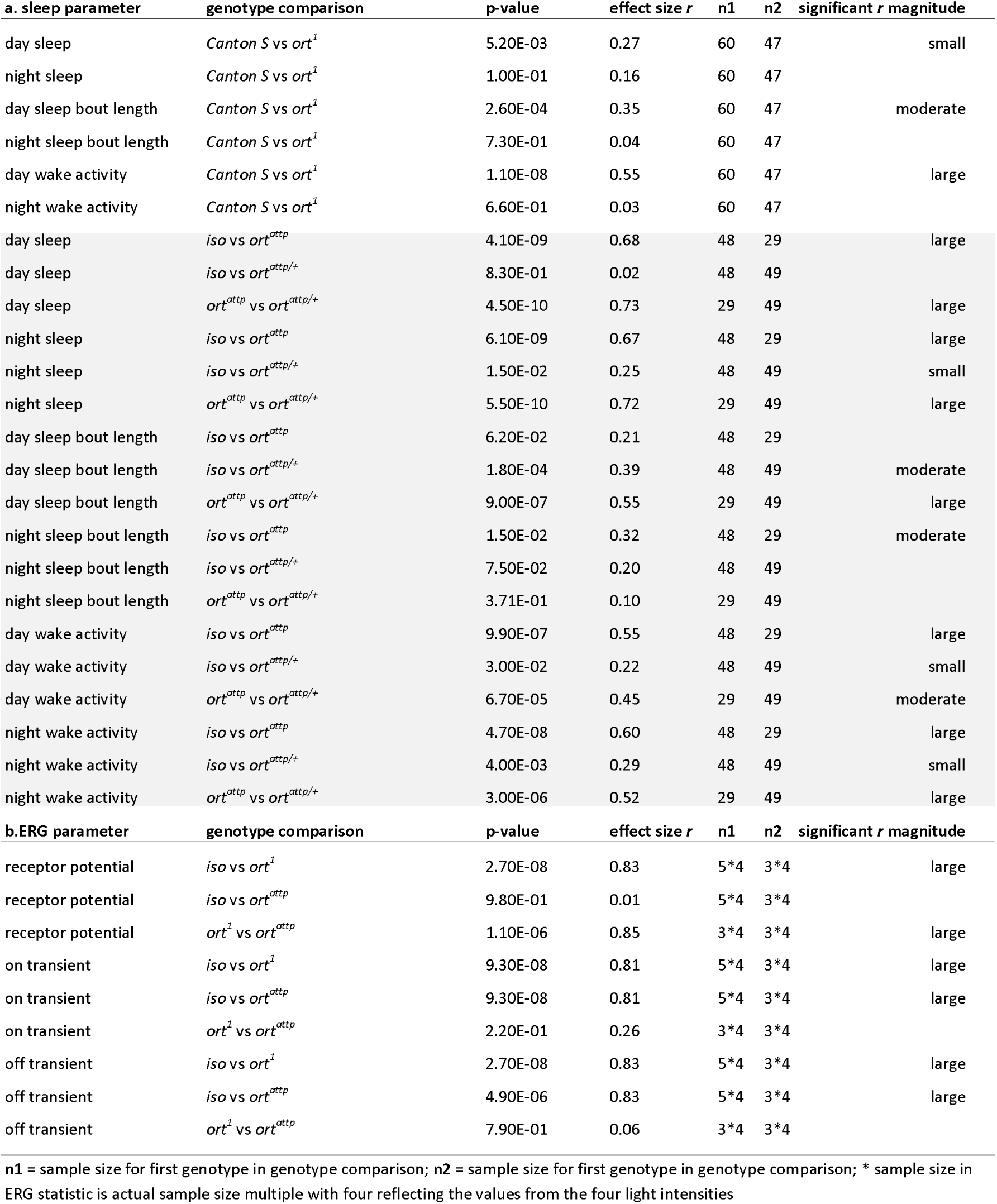
Statistics of a. sleep and b. ERG parameters for *ort* mutants.

Both *ort* mutants exhibit ERG profiles with loss of on/off transients, while *ort^1^* mutants also show loss of the receptor potential (Figure 4Hii).

### 3.3. Hyperpolarisation of Rh1-expressing photoreceptor causes short day sleep bout

Our findings so far indicate that seven of the eight phototransduction mutants exhibit day time sleep defects. However, day sleep loss in *trpl^MB^* mutants do not correlate with ERG phenotypes. Moreover, *norpA^p24^*, *trp^MB^*, *trpl^MB^*, *ort^attp/+^*, and *hdc* mutants show sleep loss in the night. Thus, we further tested if direct inhibition of photoreceptor excitability causes sleep defects in the day or night. At first, we applied the Gal4-UAS system to block synaptic transmission from all photoreceptors by expressing UAS-TeTxLC (tetanus toxin light chain) with gmr-Gal4 (as seen in Damulewicz et al., 2020). However, we were not able to obtain viable adult flies even after maintaining the culture in 22°C to reduce the known overexpression-related toxicity (Rezával et al., 2007; Sweeney et al., 1995). We next used UAS-Kir2.1 (Baines et al., 2001; Nitabach et al., 2002) crossed with gmr-Gal4 or Rh1-Gal4 to hyperpolarise all or just Rh1-expressing photoreceptors. Consistent with the finding for *ninaE* and *norpA* mutants, *gmr-Gal4>uas-Kir2.1* and *Rh1-Gal4>uas-Kir2.1* flies showed reduced day sleep as compared to *Gal4>+* or *+>uas-Kir2.1* controls (Figure 5Ai-ii, 5Di-ii), but only *Rh1-Gal4>uas-Kir2.1* flies exhibit reduced sleep bout length in the day (Figure 5Bi-ii). Night sleep parameters in both *gmr-Gal4>uas-Kir2.1* and *Rh1-Gal4>uas-Kir2.1* flies ranges between that of the controls (Figure 5E-G). This is because the unexpected low sleep exhibited in *+>uas-Kir2.1* controls (*gmr-Gal4>+* vs *+>uas-Kir2.1* and *Rh1-Gal4>+* vs *+>uas-Kir2.1*; effect size 0.62/0.74, Table 6). Therefore, we cannot determine the actual effect of hyperpolarising photoreceptor on night sleep. Kir2.1 expressing flies had no changes in waking activity (Figure 5C and 5F). Taken together, our data indicates that phototransduction and the neuronal excitability in the *Drosophila* Rh1-expressing outer photoreceptors promote day sleep.

**Figure 5.**
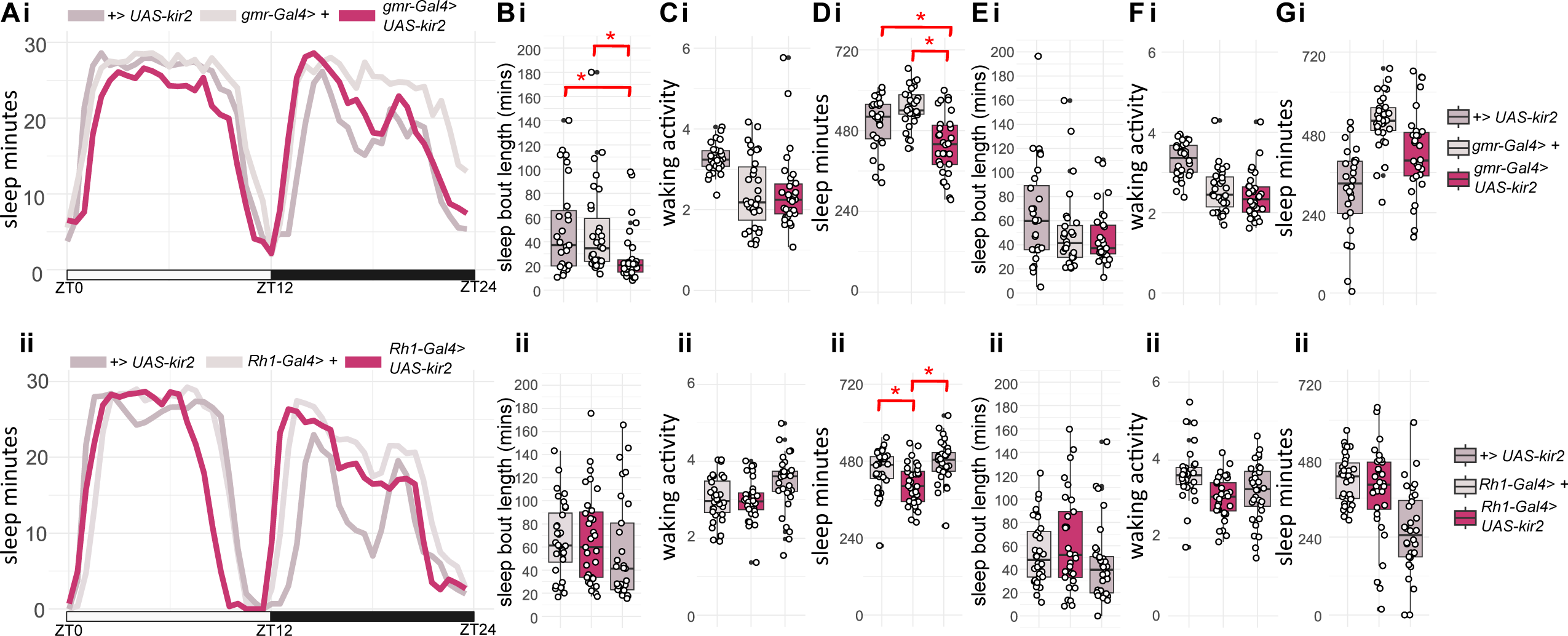
sleep profile and parameters for flies with hyperpolarised photoreceptors. **A.** Sleep traces of average sleep minutes per 30 minutes over 12 hours light-dark cycle. ZT0: light on, ZT12: light off, ZT24: end of night. Sleep traces for *gmr-Gal4>+* (i), *Rh1-Gal4>+* (ii) and *+>UAS-Kir2* (i-ii) control flies are coloured coded in the shades of greys; experimental mutants *gmr-Gal4> UAS-kir2* (i) and *Rh1-Gal4>UAS-kir2* (ii) are colour coded in light burgundy. **B.** Average day sleep bout length (minutes), **C.** Average day wake activity (beam breaks per waking minutes), **D.** Average day sleep (minutes), **E.** Average night sleep bout length (minutes), **F.** Average night wake activity (beam breaks per waking minutes), **G.** Average night sleep (minutes) for *gmr-Gal4> UAS-kir2* (i) and *Rh1-Gal4>UAS-kir2* (ii) compared to controls. Average values for each individual are plotted as filled circles except for those outside scales (B-G). Asterisk indicates statistical significance (p<0.05) between mutants and controls by pairwise Wilcoxon rank sum exact test with multiple comparison, Benjamini-Hochberg correction (followed by Kruskal-Wallis rank sum test). See **Table 6** for sample size (n=28-32), p-values and effect size for each comparison.

**Table 6.**
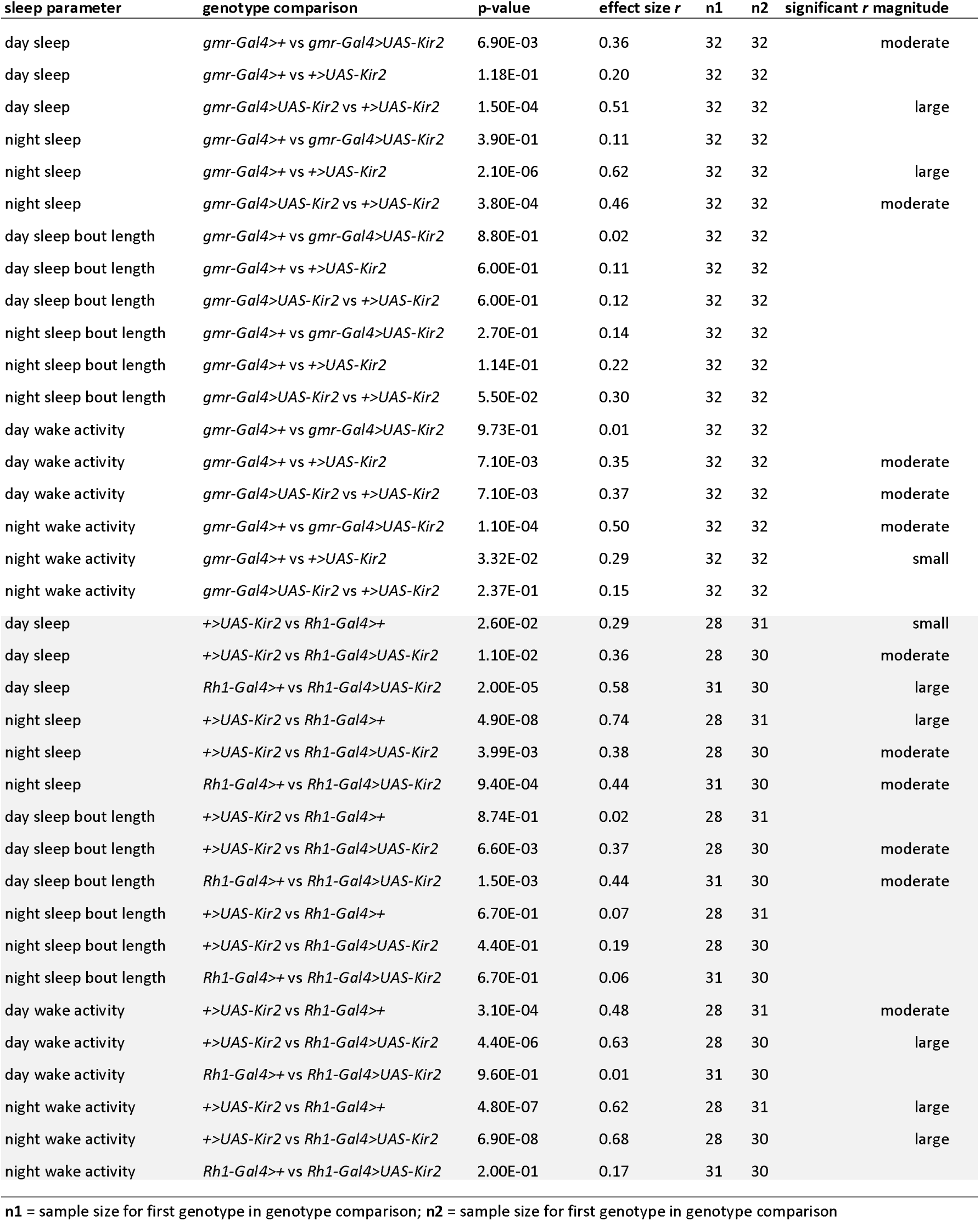
Statistics of sleep parameters for *gmrGal4>UAS-Kir2 and Rh1Gal4>UAS-Kir2* lines.

## 4. Discussion

### 4.1 Primary finding

In this study, we surveyed eight mutants with defects in phototransduction, four mutants with disrupted histaminergic transmission, the main neurotransmitter used by *Drosophila* photoreceptors, and two mutants with hyperpolarised photoreceptors. We observed short or fragmented day sleep in the majority of the phototransduction mutants as well as in mutants with hyperpolarised R1-R6 photoreceptors. This data indicates that R1-R6 photoreceptor neural activities are required for sleep homeostatic drive during the daytime. Importantly, the mutants tested in this study maintain their ability to sense light, as the non-visual pathways such as *cry*, *Rh7* and *qsm* remain intact. Nevertheless, we show robustly that reduced visual phototransduction, or downstream neurotransmission, cause sleep loss and/or sleep fragmentation, indicating a clear role for *Drosophila* photoreceptors R1-R6 in promoting day sleep.

### 4.2 Contradiction to previous studies

Our data contradicts previous findings. Specifically, it was previously shown that blocking neural transmission for R1-R6 photoreceptors via TeTxLC caused day sleep increase (Damulewicz et al., 2020). As mentioned above, we cannot produce viable adult flies that express TeTxLC in photoreceptors and instead we applied Kir2 channel to hyperpolarise retinal photoreceptors. There may be two factors that explain the data discrepancy between the two studies: Firstly, our Gal4 and UAS lines are outcrossed to the *w^iso31^* background, as is standard in the sleep research field, while such outcrossing was not described in Damulewicz et al., (2020). Secondly, vision is important for navigation, and the compromised vision in phototransduction mutants and flies with weakened photoreceptor outputs (i.e., *gmr-Gal4>UAS-kir2* or *gmr-Gal4>UAS-TeTxLC*) can result in reduced waking locomotor activities, which is a major confounding factor in assessing sleep increase. In fact, eight visual mutants in our survey (*ninaE^5^*, *ninaE^8^*, *norpA^p24^*, *trp^1^*, *trpl^302^*, *hdc^JK910^*, *ort^attp^* and *gmr-Gal4>uas-kir2*) showed significant reduced waking activity in either day and/or night. Consequentially, the reduced waking activity prevents us from concluding whether the increased night sleep seen in *ninaE^8^*, *trpl^302^* and *ort^attp^* are genuine increases or a reflection of loss of locomotion during wakefulness. Therefore, these night sleep increase phenotypes should be interpreted with caution. The waking activity was not analysed by Damulewicz et al., (2020) preventing direct comparison here.

More surprisingly, the sleep loss identified in *hdc* and *ort* mutants is also opposite to previous findings indicating that histamine is a pro-waking factor (Oh et al., 2013). One major difference between the current study and Oh et al., (2013) is that we only monitored male sleep to reach a larger range of sleep dynamics. Importantly, female flies have very low levels of day sleep (see 2.2 in Materials and method). As mentioned, further investigations of female visual mutants are necessary before it can be determined if the role of visual input to sleep diverges between male and female flies. Nevertheless, as shown in our data, two of the four mutants with histaminergic disruption show lower waking activity, consistent with the proposed arousal promoting role of histamine (Oh et al., 2013). However, we note that the mentioned previous study (Oh et al., 2013) also did not assess the impact of waking activity on sleep. Waking activity is a classic, but often ignored, analytical tool in *Drosophila* sleep research that allows differentiation between sleep and locomotor defects: as we demonstrate in this study, when the flies exhibit reduced locomotor activity, any increase in sleep cannot be certain. We therefore recommend that this parameter always be included in any locomotion-based sleep assay.

### 4.3 Experimental limitations

The unexpected abnormal low night sleep in *+>uas-Kir2* control flies which is likely due to leaky expression of Kir2 in other parts of the fly brain, since the *gmr-Gal*4 and *UAS-Kir2* lines are isogenised to *w^iso31^* background. Nevertheless, this limits our investigation to day sleep. Future investigation using gmr-LexA and LexAoP-Kir2 may be required to verify this finding.

The visual mutants tested in this study come with various genetic backgrounds including unknown *white* backgrounds and two wild type backgrounds (see Materials and methods). We therefore only interpreted the consistent phenotypes presented across the alleles. As such, there are allelic specific night sleep loss phenotypes in in *trp^MB^*, *trpl^MB^*, *ort^attp/+^* and *norpA^p24^* that is likely due to genetic background and should be interpreted with caution.

As part of control experiments, we examined the ERG profiles for all the tested gene-specific mutants. However, the severity of sleep loss phenotype does not directly correlate with ERG defects except for *trp* mutants: the weakest *ninaE^5^* mutant (only on transient reduction was detected) showed just as much day sleep fragmentation as *ninaE^8^*, *ninaE^17^* or *norpA^p24^*. Moreover, both *trpl^302^* and *trpl^MB^* exhibit sleep loss, despite minor or no impact of *trpl* mutants on ERG profile. The divergent sleep phenotypes may be partly due to the various genetic backgrounds that we did not control for (see Materials and methods). Moreover, it is important to note that ERG only reflects outer-retina physiology, not downstream optic-lobe integration. Therefore, the inconsistent sleep phenotypes can also be derived from the expression profile of phototransduction genes in the central brain beyond the photoreceptors. Future investigation utilising isogenised somatic CRISPR knockout constructs will be required to verify potential impacts of genetic background on the identified sleep-controlling role of phototransduction.

### 4.4 Hypothetical mechanisms underlying day and night sleep

Vision and light-affected sleep are mediated both via circadian clock neurons (eg., LNvs and DN1) as well as by ring neurons in the EB (Lamaze et al., 2018; Raccuglia et al., 2019; Shang et al., 2008; Singh et al., 2023). Various neural pathways connect photoreceptors to the wake promoting LNv: R1-R6 photoreceptors propagate the light signal via histamine-mediated inhibition of the downstream LMC neurons that activates LNvs (Muraro and Ceriani, 2015); on the other hand, R8 has recently been shown to communicate with LNv via the so-called AMA neurons and cholinergic signalling. Moreover, sleep promoting signals from photoreceptors may also reach the EB via the Anterior Visual Pathway through MeTu and TuBu neurons, upon which circadian clock neuron DN1 also converge (Guo et al., 2018; Lamaze et al., 2018). It is currently unclear which of the above pathways are required for the photoreceptor mediated day sleep promotion (Figure 6).

**Figure 6.**
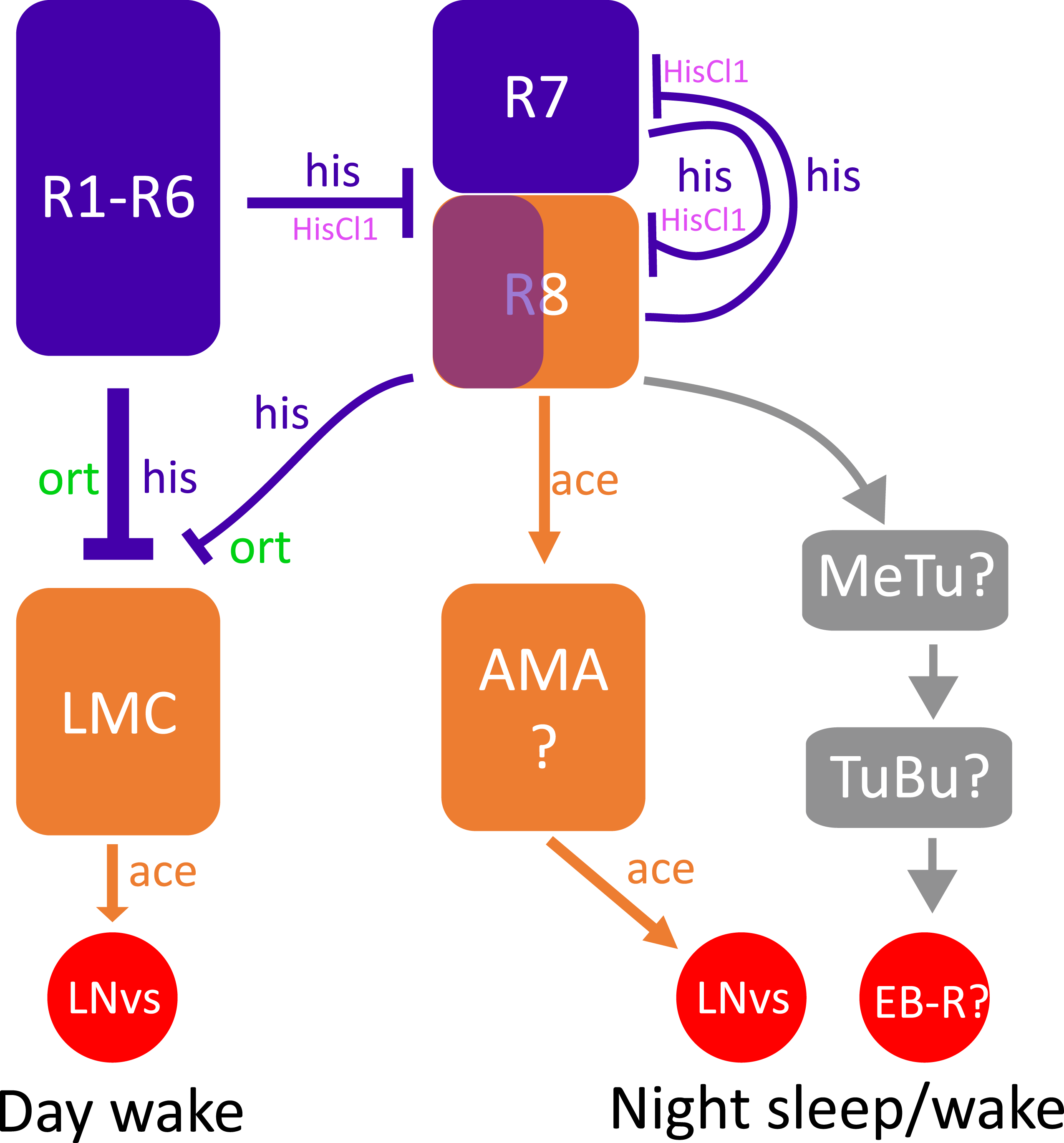
Speculative mechanism underlying visually driven day and night sleep. R1-R6 photoreceptors (blue) promote day sleep by inhibiting a subset of wake promoting lamina monopolar cells (LMC) via histamine release (his, blue text) and ort reception (green text); Night sleep is controlled by interaction between R1-R6, R7 and R8 (blue/orange) photoreceptors: Hypothetical histaminergic inhibition (his, blue text) from R1-R6 to R8 is mediated by HisCl1 (magenta text) (Alejevski et al., 2019). Separately R7 and R8 also inhibit each other via histaminergic signalling (Schnaitmann et al., 2018), these inhibitions maintain a slower suppression of acetylcholine release (ace, orange text) from R8 and its activation of the AMA neurons (orange) and downstream waking promoting LNvs (red circle). Alternatively, R8 regulate night sleep by reaching sleep-regulating EB-Ring neurons (EB-R) via MeTu and TuBu neurons with unknown neurotransmitters (grey rectangular and arrows).

Although day sleep loss in *ort^1^*, *ninaE* mutants and *Rh1-Gal4>uas-Kir2* indicates that day sleep depends on Rh1-positive R1-R6-mediated histaminergic inhibition on LMC neurons, we did not observe day sleep loss among *hdc* mutants. Nevertheless, hdc expression is wider than the visual system (Oh et al., 2013). Moreover, loss of hdc activity does not completely remove histamine production <u>(eg., Stenesen et al., 2015)</u> and it is not equal to selective blockage of ort-mediated histamine postsynaptic transmission. However, our findings are consistent with those that previously found a visually impaired *omb* mutant showed major sleep loss in both day and night (Kirszenblat et al., 2019) as well as the finding that activation of L2 LMC neurons cause day sleep lost (Muraro and Ceriani, 2015). Moreover, our data fit well with previous studies predicting that medium intensity light convey a day sleep signal via retinal photoreceptors, whereas high intensity light enhances midday sleep via HB-eyelet (Schlichting et al., 2019). Future orthogonal circuit-manipulation experimentation is required to verify if the R1-R6 photoreceptors-mediated day sleep depends on L2 LMC and/or LNvs (Figure 6).

Six tested mutants also exhibit night sleep loss (two *hdc* mutants, *norpA^p24^*, *trp^1^*, *trp^MB^* and *trpl^MB^*). Genetic background variation may contribute to this night sleep loss. However, both *hdc* mutants exhibit clear night specific sleep loss and *hdc^MB^* mutants are in an isogenic *w^iso31^* background. Therefore, this night sleep loss is likely genuine. Day time visual experience influence night sleep in flies despite there is no or limited vision in the darkness of the night. There are three possible explanations for night sleep loss in *hdc* mutants: **1.** Night sleep loss can simply be caused by non-retinal photoreceptor histaminergic neurons and is mediated by HisCl1 reception instead of ort (Oh et al., 2013). **2.** Reduction of photoreceptor-mediated histamine results in uneven inhibition among LMCs to reduce the visual contrast of bright/dark edges and differentially reduce activation of HS/VS neurons which are known promote night sleep (Kirszenblat et al., 2019). **3.** The *hdc*-mediated night sleep is related to the reduced histaminergic inhibition in non-LMC neurons mediated by HisCl1 and is independent of ort. Previous data indicates that HisCl1 is an auto-receptor for all retinal photoreceptors, and also expresses in glia cells in the optic lobe as well as clock neurons lLNvs (Davis et al., 2020; Oh et al., 2013; Pantazis et al., 2008). Among retinal photoreceptors, R8 has been proposed to receive histaminergic inhibition via HisCl1 to regulate its acetylcholine co-release (Alejevski et al., 2019; Xiao et al., 2023). Loss of this inhibition can result in an ectopic increase in acetylcholine, which can in turn activate the newly identified AMA (Xiao et al., 2023) and its downstream wake promoting lLNvs (Shang et al., 2008). Alternatively, R8 can reach sleep/wake promoting EB-R via anterior visual pathways (Figure 6).

The potential demarcation between achromatic (R1-R6) and chromatic (R8, both blue and green sensitive) photoreceptors in their control over day and night sleep (Figure 6) bears some resemblance with the previous finding in mammalian system, where the sleep- vs wake-promoting effects of green and blue light required proper neural output from rod/cone and the melanopsin expressing photosensitive retinal ganglion cells (Pilorz et al., 2016). Nevertheless, the revelation of the potential neural network demarcation of day and night sleep in *Drosophila* require further investigation to expand our understandings in circuital logic of light-mediated sleep regulation.

## Acknowledgements

This study was funded by University of Leicester to M.A. and X.L. (F50/100 PhD studentships), N. S., S. H., T. O., and J. A (PGT and UG programme funding), an Erasmus + studentship to C. S.-G., a Royal Society University Fellowship to B. W., and a BBSRC New Investigator Scheme (BB/W014939/1) to K-F.C and Y-C. H. We thank Dr. Kalina Davies for her proofreading.

## Author Contributions

K-F.C – Conceptualization, Methodology, Investigation, Writing - Original Draft, Writing – Review & Editing, Visualization, Supervision, Project administration, Funding acquisition; Y.-C. H. – Investigation, Visualization, Writing – Review & Editing. B.W. – Methodology, Investigation, Writing – Review & Editing. M.A., X.L., N. S., S. H., T. O., J. A, and C. S.-G., – Investigation, Writing – Review & Editing.

## Reference

Alejevski, F., Saint-Charles, A., Michard-Vanhee, C., Martin, B., Galant, S., Vasiliauskas, D., Rouyer, F., 2019. The HisCl1 histamine receptor acts in photoreceptors to synchronize Drosophila behavioral rhythms with light-dark cycles. Nature communications 10, 252. 10.1038/s41467-018-08116-7

Alphen, B. van, Semenza, E.R., Yap, M., Swinderen, B. van, Allada, R., 2021. A deep sleep stage in Drosophila with a functional role in waste clearance. Science Advances 7, eabc2999. 10.1126/sciadv.abc2999

Altimus, C.M., Güler, A.D., Villa, K.L., McNeill, D.S., LeGates, T.A., Hattar, S., 2008. Rods-cones and melanopsin detect light and dark to modulate sleep independent of image formation. Proceedings of the National Academy of Sciences 105, 19998–20003. 10.1073/pnas.0808312105

Andretic, R., Shaw, P.J., 2005. Essentials of Sleep Recordings in *Drosophila:* Moving Beyond Sleep Time, in: Young, M.W. (Ed.), Methods in Enzymology, Circadian Rhythms. Academic Press, pp. 759–772. 10.1016/S0076-6879(05)93040-1

Baines, R.A., Uhler, J.P., Thompson, A., Sweeney, S.T., Bate, M., 2001. Altered Electrical Properties in DrosophilaNeurons Developing without Synaptic Transmission. J. Neurosci. 21, 1523–1531. 10.1523/JNEUROSCI.21-05-01523.2001

Burg, M.G., Sarthy, P.V., Koliantz, G., Pak, W.L., 1993. Genetic and molecular identification of a Drosophila histidine decarboxylase gene required in photoreceptor transmitter synthesis. The EMBO Journal 12, 911–919. 10.1002/j.1460-2075.1993.tb05732.x

Chen, K.F., Lowe, S., Lamaze, A., Kratschmer, P., Jepson, J., 2019. Neurocalcin regulates nighttime sleep and arousal in Drosophila. eLife 8. 10.7554/eLife.38114

Chen, K.F., Peschel, N., Zavodska, R., Sehadova, H., Stanewsky, R., 2011. QUASIMODO, a Novel GPI-anchored zona pellucida protein involved in light input to the Drosophila circadian clock. Current Biology 21, 719–729.

Chen, Z., Chen, H.-C., Montell, C., 2015. TRP and Rhodopsin Transport Depends on Dual XPORT ER Chaperones Encoded by an Operon. Cell Reports 13, 573–584. 10.1016/j.celrep.2015.09.018

Chiu, J.C., Low, K.H., Pike, D.H., Yildirim, E., Edery, I., 2010. Assaying Locomotor Activity to Study Circadian Rhythms and Sleep Parameters in Drosophila. JoVE (Journal of Visualized Experiments) e2157. 10.3791/2157

Cosens, D.J., Manning, A., 1969. Abnormal Electroretinogram from a Drosophila Mutant. Nature 224, 285–287. 10.1038/224285a0

Damulewicz, M., Ispizua, J.I., Ceriani, M.F., Pyza, E.M., 2020. Communication Among Photoreceptors and the Central Clock Affects Sleep Profile. Front. Physiol. 11. 10.3389/fphys.2020.00993

Davis, F.P., Nern, A., Picard, S., Reiser, M.B., Rubin, G.M., Eddy, S.R., Henry, G.L., 2020. A genetic, genomic, and computational resource for exploring neural circuit function. eLife 9, e50901. 10.7554/eLife.50901

Deng, B., Li, Q., Liu, X., Cao, Y., Li, B., Qian, Y., Xu, R., Mao, R., Zhou, E., Zhang, W., Huang, J., Rao, Y., 2019. Chemoconnectomics: Mapping Chemical Transmission in Drosophila. Neuron 101, 876–893.e4. 10.1016/j.neuron.2019.01.045

Dissel, S., Angadi, V., Kirszenblat, L., Suzuki, Y., Donlea, J., Klose, M., Koch, Z., English, D., Winsky-Sommerer, R., van Swinderen, B., Shaw, P.J., 2015. Sleep Restores Behavioral Plasticity to Drosophila Mutants. Current Biology 25, 1270–1281. 10.1016/j.cub.2015.03.027

Dove, A.E., Cook, B.L., Irgebay, Z., Vecsey, C.G., 2017. Mechanisms of sleep plasticity due to sexual experience in *Drosophila melanogaster*. Physiology & Behavior 180, 146–158. 10.1016/j.physbeh.2017.08.020

Duhart, J.M., Buchler, J.R., Inami, S., Kennedy, K.J., Jenny, B.P., Afonso, D.J.S., Koh, K., 2023. Modulation and neural correlates of postmating sleep plasticity in Drosophila females. Current Biology 33, 2702–2716.e3. 10.1016/j.cub.2023.05.054

Gao, S., Takemura, S., Ting, C.-Y., Huang, S., Lu, Z., Luan, H., Rister, J., Thum, A.S., Yang, M., Hong, S.-T., Wang, J.W., Odenwald, W.F., White, B.H., Meinertzhagen, I.A., Lee, C.-H., 2008. The Neural Substrate of Spectral Preference in Drosophila. Neuron 60, 328–342. 10.1016/j.neuron.2008.08.010

Gilestro, G.F., Tononi, G., Cirelli, C., 2009. Widespread Changes in Synaptic Markers as a Function of Sleep and Wakefulness in Drosophila. Science 324, 109–112. 10.1126/science.1166673

Guo, F., Holla, M., Díaz, M.M., Rosbash, M., 2018. A Circadian Output Circuit Controls Sleep-Wake Arousal in Drosophila. Neuron 100, 624–635.e4. 10.1016/j.neuron.2018.09.002

Harbison, S.T., Sehgal, A., 2008. Quantitative Genetic Analysis of Sleep in Drosophila melanogaster. Genetics 178, 2341–2360. 10.1534/genetics.107.081232

Hardie, R.C., 2012. Phototransduction mechanisms in Drosophila microvillar photoreceptors. Wiley Interdisciplinary Reviews: Membrane Transport and Signaling 1, 162–187. 10.1002/wmts.20

Hartmann, C., Kempf, A., 2023. Mitochondrial control of sleep. Current Opinion in Neurobiology 81, 102733. 10.1016/j.conb.2023.102733

Horne, J.A., Walmsley, B., 1976. Daytime visual load and the effects upon human sleep. Psychophysiology 13, 115–20.

Hsu, C.T., Guevara, C., Killiany, S.L., Shon, J., Dissel, S., Sehgal, A., 2025. Clock-dependent regulation of a homeostatic sleep center maintains daytime sleep and evening activity. Current Biology 0. 10.1016/j.cub.2025.06.003

Inoue, H., Yoshioka, T., Hotta, Y., 1989. Diacylglycerol Kinase Defect in a *Drosophila* Retinal Degeneration Mutant *rdgA*. Journal of Biological Chemistry 264, 5996–6000. 10.1016/S0021-9258(18)83648-1

Isaac, R.E., Li, C., Leedale, A.E., Shirras, A.D., 2009. Drosophila male sex peptide inhibits siesta sleep and promotes locomotor activity in the post-mated female. Proceedings of the Royal Society of London B: Biological Sciences rspb20091236.

Ishimoto, H., Lark, A., Kitamoto, T., 2012. Factors that differentially affect daytime and nighttime sleep in Drosophila melanogaster. Front Neurol 3, 24.

Jagannathan, S.R., Jeans, T., Van De Poll, M.N., van Swinderen, B., 2024. Multivariate classification of multichannel long-term electrophysiology data identifies different sleep stages in fruit flies. Science Advances 10, eadj4399. 10.1126/sciadv.adj4399

Jones, J.D., Holder, B.L., Montgomery, A.C., McAdams, C.V., He, E., Burns, A.E., Eiken, K.R., Vogt, A., Velarde, A.I., Elder, A.J., McEllin, J.A., Dissel, S., 2025. The dorsal fan-shaped body is a neurochemically heterogeneous sleep-regulating center in Drosophila. PLOS Biology 23, e3003014. 10.1371/journal.pbio.3003014

Kirszenblat, L., Yaun, R., van Swinderen, B., 2019. Visual experience drives sleep need in Drosophila. Sleep 42. 10.1093/sleep/zsz102

Korf, E.M., Mölle, M., Born, J., Ngo, H.-V.V., 2017. Blindfolding during wakefulness causes decrease in sleep slow wave activity. Physiological Reports 5, e13239. 10.14814/phy2.13239

Lakhiani, R., Shanavas, S., Melnattur, K., 2023. Comparative biology of sleep in diverse animals. J Exp Biol 226. 10.1242/jeb.245677

Lamaze, A., Krätschmer, P., Chen, K.-F., Lowe, S., Jepson, J.E.C., 2018. A Wake-Promoting Circadian Output Circuit in Drosophila. Current Biology 28, 3098–3105.e3. 10.1016/j.cub.2018.07.024

Leung, H.-T., Geng, C., Pak, W.L., 2000. Phenotypes of trpl Mutants and Interactions between the Transient Receptor Potential (TRP) and TRP-Like Channels inDrosophila. J. Neurosci. 20, 6797–6803. 10.1523/JNEUROSCI.20-18-06797.2000

Mazzotta, G.M., Damulewicz, M., Cusumano, P., 2020. Better Sleep at Night: How Light Influences Sleep in Drosophila. Front. Physiol. 11. 10.3389/fphys.2020.00997

Meyerhof, G.T., Easwaran, S., Bontempo, A.E., Montell, C., Montell, D.J., 2024. Altered circadian rhythm, sleep, and rhodopsin 7–dependent shade preference during diapause in Drosophila melanogaster. Proceedings of the National Academy of Sciences 121, e2400964121. 10.1073/pnas.2400964121

Miyamoto, H., Katagiri, H., Hensch, T., 2003. Experience-dependent slow-wave sleep development. Nat Neurosci 6, 553–554. 10.1038/nn1064

Muraro, N.I., Ceriani, M.F., 2015. Acetylcholine from visual circuits modulates the activity of arousal neurons in Drosophila. Journal of Neuroscience 35, 16315–16327.

Nériec, N., Desplan, C., 2016. Chapter Fourteen - From the Eye to the Brain: Development of the Drosophila Visual System, in: Wassarman, P.M. (Ed.), Current Topics in Developmental Biology, Essays on Developmental Biology, Part A. Academic Press, pp. 247–271. 10.1016/bs.ctdb.2015.11.032

Ni, J.D., Baik, L.S., Holmes, T.C., Montell, C., 2017. A rhodopsin in the brain functions in circadian photoentrainment in Drosophila. Nature 545, 340–344. 10.1038/nature22325

Niemeyer, B.A., Suzuki, E., Scott, K., Jalink, K., Zuker, C.S., 1996. The Drosophila Light-Activated Conductance Is Composed of the Two Channels TRP and TRPL. Cell 85, 651–659. 10.1016/S0092-8674(00)81232-5

Nitabach, M.N., Blau, J., Holmes, T.C., 2002. Electrical Silencing of Drosophila Pacemaker Neurons Stops the Free-Running Circadian Clock. Cell 109, 485–495. 10.1016/S0092-8674(02)00737-7

Ogueta, M., Hardie, R.C., Stanewsky, R., 2020. Light Sampling via Throttled Visual Phototransduction Robustly Synchronizes the Drosophila Circadian Clock. Current Biology 30, 2551–2563.e3. 10.1016/j.cub.2020.04.067

Ogueta, M., Hardie, R.C., Stanewsky, R., 2018. Non-canonical Phototransduction Mediates Synchronization of the Drosophila melanogaster Circadian Clock and Retinal Light Responses. Current Biology 28, 1725–1735.e3. 10.1016/j.cub.2018.04.016

Oh, Y., Jang, D., Sonn, J.Y., Choe, J., 2013. Histamine-HisCl1 Receptor Axis Regulates Wake-Promoting Signals in Drosophila melanogaster. PLoS ONE 8, e68269. 10.1371/journal.pone.0068269

O’Tousa, J.E., Baehr, W., Martin, R.L., Hirsh, J., Pak, W.L., Applebury, M.L., 1985. The Drosophila *ninaE* gene encodes an opsin. Cell 40, 839–850. 10.1016/0092-8674(85)90343-5

Pantazis, A., Segaran, A., Liu, C.-H., Nikolaev, A., Rister, J., Thum, A.S., Roeder, T., Semenov, E., Juusola, M., Hardie, R.C., 2008. Distinct Roles for Two Histamine Receptors (hclA and hclB) at the Drosophila Photoreceptor Synapse. The Journal of Neuroscience 28, 7250–7259. 10.1523/jneurosci.1654-08.2008

Peschel, N., Chen, K.F., Szabo, G., Stanewsky, R., 2009. Light-dependent interactions between the Drosophila circadian clock factors cryptochrome, jetlag, and timeless. Current Biology 19, 241–247.

Pilorz, V., Tam, S.K.E., Hughes, S., Pothecary, C.A., Jagannath, A., Hankins, M.W., Bannerman, D.M., Lightman, S.L., Vyazovskiy, V.V., Nolan, P.M., Foster, R.G., Peirson, S.N., 2016. Melanopsin Regulates Both Sleep-Promoting and Arousal-Promoting Responses to Light. PLOS Biology 14, e1002482. 10.1371/journal.pbio.1002482

Raccuglia, D., Huang, S., Ender, A., Heim, M.M., Laber, D., Suárez-Grimalt, R., Liotta, A., Sigrist, S.J., Geiger, J.R.P., Owald, D., 2019. Network-Specific Synchronization of Electrical Slow-Wave Oscillations Regulates Sleep Drive in Drosophila. Current Biology. 10.1016/j.cub.2019.08.070

Reinhard, N., Fukuda, A., Manoli, G., Derksen, E., Saito, A., Möller, G., Sekiguchi, M., Rieger, D., Helfrich-Förster, C., Yoshii, T., Zandawala, M., 2024. Synaptic connectome of the Drosophila circadian clock. Nat Commun 15, 10392. 10.1038/s41467-024-54694-0

Rezával, C., Werbajh, S., Ceriani, M.F., 2007. Neuronal death in Drosophila triggered by GAL4 accumulation. European Journal of Neuroscience 25, 683–694. 10.1111/j.1460-9568.2007.05317.x

Rieger, D., Fraunholz, C., Popp, J., Bichler, D., Dittmann, R., Helfrich-Förster, C., 2007. The Fruit Fly Drosophila melanogaster Favors Dim Light and Times Its Activity Peaks to Early Dawn and Late Dusk. J Biol Rhythms 22, 387–399. 10.1177/0748730407306198

Rieger, D., Stanewsky, R., Helfrich-Förster, C., 2003. Cryptochrome, Compound Eyes, Hofbauer-Buchner Eyelets, and Ocelli Play Different Roles in the Entrainment and Masking Pathway of the Locomotor Activity Rhythm in the Fruit Fly Drosophila Melanogaster. Journal of Biological Rhythms 18, 377–391. 10.1177/0748730403256997

Rossi, F.A., Bochicchio, P.A., Quesada-Allué, L.A., Pérez, M.M., 2015. N-β-alanyldopamine metabolism, locomotor activity and sleep in Drosophila melanogaster ebony and tan mutants. Physiological Entomology 40, 166–174. 10.1111/phen.12100

Scavarda, N.J., O’Tousa, J., Pak, W.L., 1983. Drosophila locus with gene-dosage effects on rhodopsin. Proceedings of the National Academy of Sciences 80, 4441–4445. 10.1073/pnas.80.14.4441

Schlichting, M., Menegazzi, P., Lelito, K.R., Yao, Z., Buhl, E., Benetta, E.D., Bahle, A., Denike, J., Hodge, J.J., Helfrich-Förster, C., Shafer, O.T., 2016. A Neural Network Underlying Circadian Entrainment and Photoperiodic Adjustment of Sleep and Activity in Drosophila. J. Neurosci. 36, 9084–9096. 10.1523/JNEUROSCI.0992-16.2016

Schlichting, M., Menegazzi, P., Rosbash, M., Helfrich-Förster, C., 2019. A Distinct Visual Pathway Mediates High-Intensity Light Adaptation of the Circadian Clock in Drosophila. J. Neurosci. 39, 1621–1630. 10.1523/JNEUROSCI.1497-18.2018

Schnaitmann, C., Haikala, V., Abraham, E., Oberhauser, V., Thestrup, T., Griesbeck, O., Reiff, D.F., 2018. Color Processing in the Early Visual System of Drosophila. Cell 172, 318–330.e18. 10.1016/j.cell.2017.12.018

Sengupta, S., Crowe, L.B., You, S., Roberts, M.A., Jackson, F.R., 2019. A Secreted Ig-Domain Protein Required in Both Astrocytes and Neurons for Regulation of *Drosophila* Night Sleep. Current Biology 29, 2547–2554.e2. 10.1016/j.cub.2019.06.055

Shafer, O.T., Keene, A.C., 2021. The Regulation of Drosophila Sleep. Current Biology 31, R38–R49. 10.1016/j.cub.2020.10.082

Shang, Y., Griffith, L.C., Rosbash, M., 2008. Light-arousal and circadian photoreception circuits intersect at the large PDF cells of the *Drosophila* brain. Proceedings of the National Academy of Sciences 105, 19587–19594. 10.1073/pnas.0809577105

Singh, P., Aleman, A., Omoto, J.J., Nguyen, B.-C., Kandimalla, P., Hartenstein, V., Donlea, J.M., 2023. Examining Sleep Modulation by Drosophila Ellipsoid Body Neurons. eNeuro 10. 10.1523/ENEURO.0281-23.2023

Stenesen, D., Moehlman, A.T., Krämer, H., 2015. The carcinine transporter CarT is required in Drosophila photoreceptor neurons to sustain histamine recycling. eLife 4, e10972. 10.7554/eLife.10972

Suh, J., Jackson, F.R., 2007. Drosophila Ebony Activity Is Required in Glia for the Circadian Regulation of Locomotor Activity. Neuron 55, 435–447. 10.1016/j.neuron.2007.06.038

Sweeney, S.T., Broadie, K., Keane, J., Niemann, H., O’Kane, C.J., 1995. Targeted expression of tetanus toxin light chain in Drosophila specifically eliminates synaptic transmission and causes behavioral defects. Neuron 14, 341–351. 10.1016/0896-6273(95)90290-2

Tainton-Heap, L.A.L., Kirszenblat, L.C., Notaras, E.T., Grabowska, M.J., Jeans, R., Feng, K., Shaw, P.J., van Swinderen, B., 2021. A Paradoxical Kind of Sleep in Drosophila melanogaster. Current Biology 31, 578–590.e6. 10.1016/j.cub.2020.10.081

Tang, M., Cao, L.-H., Yang, T., Ma, S.-X., Jing, B.-Y., Xiao, N., Xu, S., Leng, K.-R., Yang, D., Li, M.-T., Luo, D.-G., 2022. An extra-clock ultradian brain oscillator sustains circadian timekeeping. Science Advances 8, eabo5506. 10.1126/sciadv.abo5506

Vaccaro, A., Dor, Y.K., Nambara, K., Pollina, E.A., Lin, C., Greenberg, M.E., Rogulja, D., 2020. Sleep Loss Can Cause Death through Accumulation of Reactive Oxygen Species in the Gut. Cell 0. 10.1016/j.cell.2020.04.049

Vincent, C.M., Beckwith, E.J., Silva, C.J.S. da, Pearson, W.H., Kierdorf, K., Gilestro, G.F., Dionne, M.S., 2022. Infection increases activity via Toll dependent and independent mechanisms in Drosophila melanogaster. PLOS Pathogens 18, e1010826. 10.1371/journal.ppat.1010826

Wu, B., Ma, L., Zhang, E., Du, J., Liu, S., Price, J., Li, S., Zhao, Z., 2018. Sexual dimorphism of sleep regulated by juvenile hormone signaling in Drosophila. PLOS Genetics 14, e1007318. 10.1371/journal.pgen.1007318

Wulff, K., Gatti, S., Wettstein, J.G., Foster, R.G., 2010. Sleep and circadian rhythm disruption in psychiatric and neurodegenerative disease. Nature Reviews Neuroscience 11, 589–599. 10.1038/nrn2868

Xiao, N., Xu, S., Li, Z.-K., Tang, M., Mao, R., Yang, T., Ma, S.-X., Wang, P.-H., Li, M.-T., Sunilkumar, A., Rouyer, F., Cao, L.-H., Luo, D.-G., 2023. A single photoreceptor splits perception and entrainment by cotransmission. Nature 623, 562–570. 10.1038/s41586-023-06681-6

Yap, M.H.W., Grabowska, M.J., Rohrscheib, C., Jeans, R., Troup, M., Paulk, A.C., van Alphen, B., Shaw, P.J., van Swinderen, B., 2017. Oscillatory brain activity in spontaneous and induced sleep stages in flies. Nat Commun 8, 1815. 10.1038/s41467-017-02024-y

Zanini, D., Giraldo, D., Warren, B., Katana, R., Andrés, M., Reddy, S., Pauls, S., Schwedhelm-Domeyer, N., Geurten, B.R.H., Göpfert, M.C., 2018. Proprioceptive Opsin Functions in Drosophila Larval Locomotion. Neuron 98, 67–74.e4. 10.1016/j.neuron.2018.02.028

